# Archaeal type six secretion system mediates contact-dependent antagonism

**DOI:** 10.1101/2024.04.11.588991

**Authors:** Tobias Zachs, Jessie James L. Malit, Jingwei Xu, Alexandra Schürch, Shamphavi Sivabalasarma, Phillip Nußbaum, Sonja-Verena Albers, Martin Pilhofer

**Affiliations:** Department of Biology, Institute of Molecular Biology & Biophysics, Eidgenössische Technische Hochschule Zürich, Otto-Stern-Weg 5, 8093 Zürich, Switzerland; Molecular Biology of Archaea, Institute of Biology, Faculty of Biology, University of Freiburg, Schänzlestr. 1, 79104 Freiburg, Germany; Spemann Graduate School of Biology and Medicine, University of Freiburg, 79104 Freiburg, Germany

## Abstract

Microbial communities are shaped by cell-cell interactions. Even though archaea are often found in associations with other microorganisms, the mechanisms structuring these communities are poorly understood. Here we report the structure and function of haloarchaeal contractile injection systems (CISs). Using a combination of functional assays and time lapse imaging, we show that Halogeometricum borinquense exhibits antagonism towards Haloferax volcanii by inducing cell lysis and inhibiting proliferation. This antagonism is contact-dependent and requires a functional CIS, which is encoded by a gene cluster that is associated with toxin-immunity pairs. Cryo-focused ion beam milling and imaging by cryo-electron tomography revealed CISs bound to the cytoplasmic membrane, resembling bacterial type six secretion systems (T6SSs). We show that related T6SS gene clusters are conserved and expressed in other haloarchaeal strains with antagonistic behavior. Our data provides a mechanistic framework for understanding how archaea may shape microbial communities and impact the food webs they inhabit.

**Teaser:** T6SSs are widespread in the archaeal domain and used to kill other archaea.

## Introduction

Ecological niches frequently contain a complex, yet defined, mixture of different microorganisms (*1*, *2*). Bacteria employ a wide range of highly adapted secretion systems to translocate arsenals of toxins across their cell envelopes (*3*). This allows bacteria to obtain a competitive advantage by structuring their surrounding community and also grants them access to alternative substrate sources. Archaea are also known to exist in complex microbiomes, including biofilms and microbial consortia (*2*, *4*). Studies of these communities have primarily focused on the metabolic flux between different members (*5–8*). With the exception of archaeocins, mechanistic insights into archaeal antagonism and how antagonism is used to define an archaeal niche is largely lacking (*9–11*).

Interestingly, recent bioinformatic studies revealed the presence of contractile injection system (CIS) gene clusters in archaeal genomes from different phyla (*11–14*). Related CIS gene clusters are widespread among diverse bacteria and known to mediate a wide range of interactions between bacteria and other organisms (*3*). With a structure and mechanism that is evolutionarily related to contractile phages, like the T4 phage, CISs allow the host to translocate effector proteins into the surrounding environment or directly into neighboring cells (*15–17*). CISs have adapted to a range of functions, but their conserved core is composed of a sheath-tube module (also called a contractile tail), which assembles onto a baseplate. To facilitate toxin secretion, the baseplate triggers contraction of a dynamic outer sheath, forcing a rigid inner tube, tipped with a spike protein, to be driven through membranes (*18*). This dynamic transition from a primed extended state to a fired contracted state is a hallmark of CISs. The process by which this occurs – i.e. their mode-of-action – can vary.

The two most commonly studied modes-of-action are the “type six secretion systems” (T6SSs) and the “extracellular CISs” (eCISs). T6SSs assemble and fire while attached to the host’s cytoplasmic membrane. To date, T6SSs have only been described in Gram-negative bacteria, which use these systems to translocate cargo across the cell envelope in a cell-cell contact-dependent manner (*19*, *20*). eCISs, in contrast, are assembled within the bacterial host cell but are only functional when released into the environment by cell lysis. Once in contact with a target cell, eCIS use their tail fibers to bind and subsequently contract to inject their cargo (*21–24*).

The bioinformatic identification of CIS gene clusters in archaea raises pressing questions regarding their expression, mode-of-action and function; offering exciting prospects for understanding how archaea may structure the communities that they thrive in.

## Results

### *H. borinquense* assembles contractile sheaths composed of Cis2

We chose to establish *Halogeometricum borinquense* (hereafter *H. borinquense* or *Hb*) as the model system for investigating archaeal CIS gene clusters. *Hb* is a halophilic archaeon that was originally isolated from the solar salterns of Cabo Rojo, Puerto Rico (*25*, *26*). Its genome was reported to possess genes with similarities to a diverse group of related CISs (*13*). These CIS gene clusters are phylogenetically part of “clade IIa”, which to date has no characterized representatives. We re-analyzed the *Hb* CIS gene cluster (accessions: *Hbor_38640-38890*), which is encoded on a 210 kb genomic plasmid (pHBOR03) (Figure 1A). 14 genes with similarities to previously identified structural components of bacterial CISs are present in the gene cluster. These structural genes are organized in two sub-clusters, which are separated by an ‘interspacing’ region of ∼8.7 kb, containing genes for nine hypothetical proteins. The 14 structural genes were re-named according to their homologies to previously characterized components of other CISs (Table 1) (*24*, *27*). All but homologs for the plug (*cis6*), tape measure (*cis14*) and crown (*cis19*) were found in the gene cluster.

**Figure 1:**
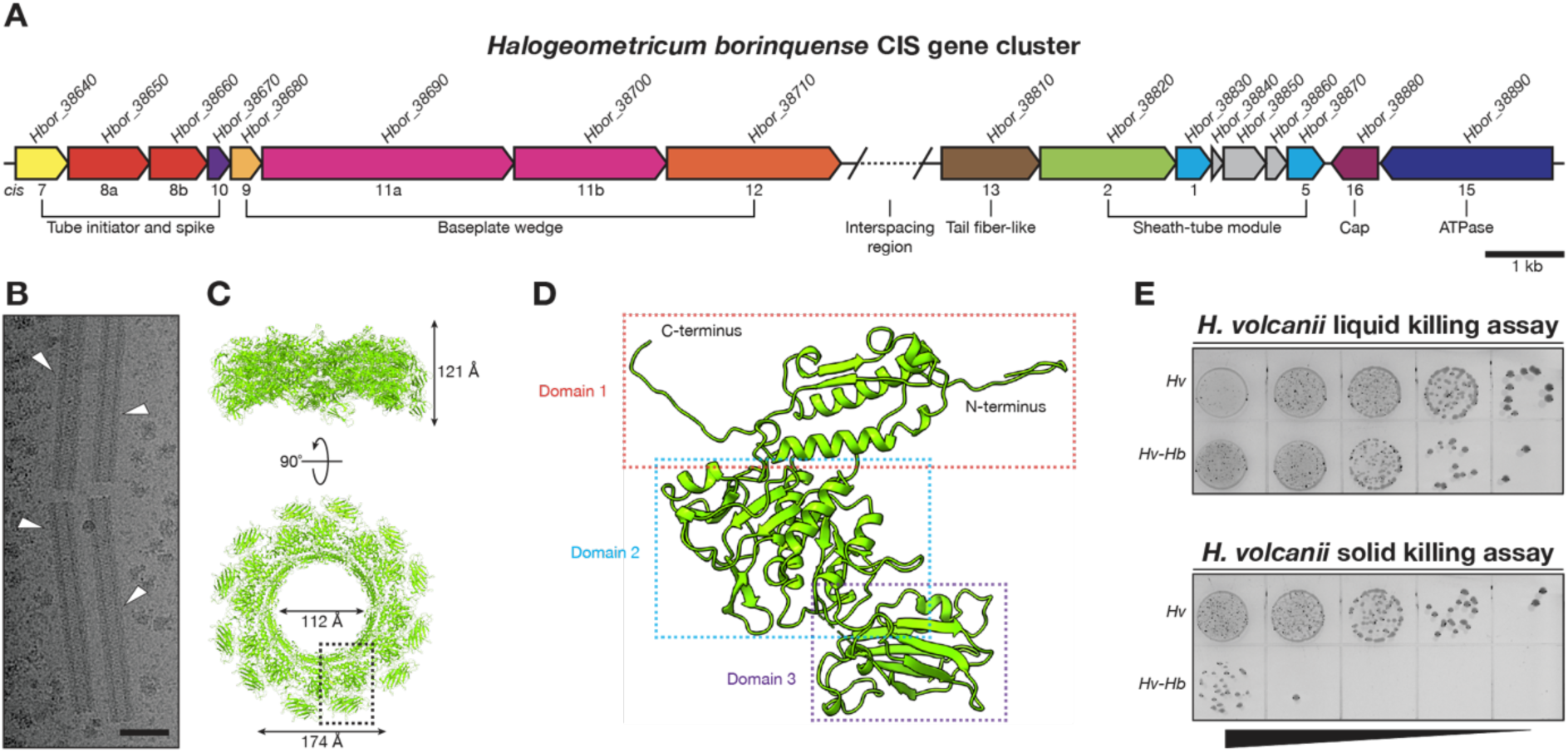
*H. borinquense* expresses CIS genes and exhibits antagonism. **(A)** Schematic of the *H. borinquense* CIS gene cluster found on the plasmid pHBOR03. Annotations include the gene accession numbers (*Hbor_28640-38890*) and the CIS gene nomenclature (*cis1-16*) with genes grouped together by predicted functional groups. For details on the interspacing region see Figure S4. **(B)** Micrograph of plunge frozen sheath isolated from *H. borinquense*. Micrograph shows sheath structures used for single particle helical reconstruction (white arrowheads). Sheath lengths were heterogeneous. Scale bar: 50 nm. **(C)** Side and top views of four layers of the contracted sheath structure composed of Cis2 with corresponding dimensions. Single subunit within dashed box is depicted in panel **D**. **(D)** Structure of the contracted sheath monomer derived from a single particle helical reconstruction at a resolution of 2.8 Å. Within the sheath monomer, three domains are observable with only a section between domain 2 and 3 being unresolved due to flexibility (residues 209-291). **(E)** Killing assay performed by co-incubating *H. borinquense* (*Hb*) with *H. volcanii* (*Hv*) in liquid and on solid media for 24 h at 45°C. Shown are serial dilutions spots (10^−2^ to 10^−6^) spotted on a *H. volcanii* selection plate. Note the significant drop (1,000-fold) in the growth of *H. volcanii* at higher dilutions in the solid killing assay.

**Table 1:**
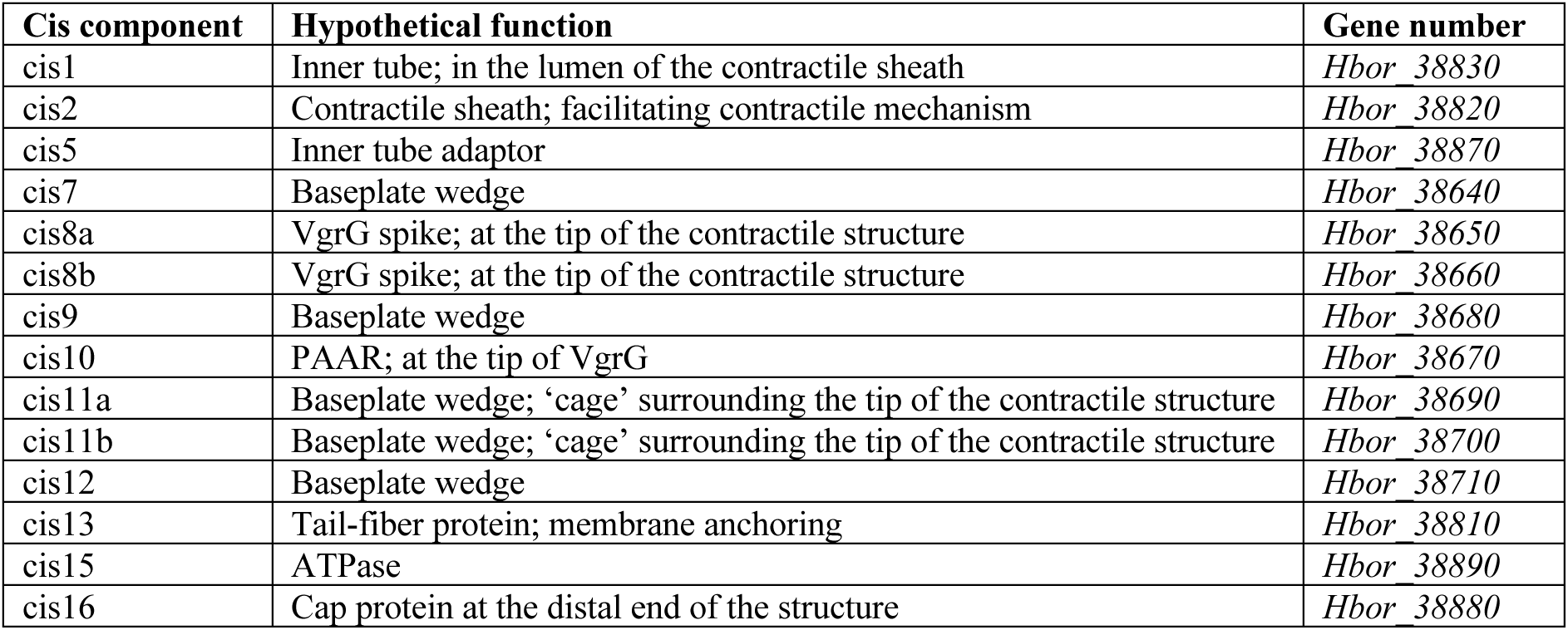
*H. borinquense* cis components.

| **Cis component** | **Hypothetical function** | **Gene number** |
| --- | --- | --- |
| cis1 | Inner tube; in the lumen of the contractile sheath | *Hbor* 38830 |
| cis2 | Contractile sheath; facilitating contractile mechanism | *Hbor* 38820 |
| cis5 | Inner tube adaptor | *Hbor* 38870 |
| cis7 | Baseplate wedge | *Hbor* 38640 |
| cis8a | VgrG spike; at the tip of the contractile structure | *Hbor* 38650 |
| cis8b | VgrG spike; at the tip of the contractile structure | *Hbor* 38660 |
| cis9 | Baseplate wedge | *Hbor* 38680 |
| cis10 | PAAR; at the tip of VgrG | *Hbor* 38670 |
| cis11a | Baseplate wedge; 'cage' surrounding the tip of the contractile structure | *Hbor* 38690 |
| cis11b | Baseplate wedge; 'cage' surrounding the tip of the contractile structure | *Hbor* 38700 |
| cis12 | Baseplate wedge | *Hbor* 38710 |
| cis13 | Tail-fiber protein; membrane anchoring | *Hbor* 38810 |
| cis15 | ATPase | *Hbor* 38890 |
| cis16 | Cap protein at the distal end of the structure | *Hbor* 38880 |

Next, we tested the expression of *cis* genes by performing crude purifications of CISs from *Hb* cultures. Electron microscopy (EM) imaging revealed an abundance of sheath-like structures that were always in a contracted state with a mean length of 220 ±16 nm (Figure 1B; Figure S1A/B). Mass spectrometry analysis of the preparations detected the presence of *Hb* protein Cis2 (*Hbor_38820*), but no other CIS gene cluster components were identified. Using single particle cryoEM and helical reconstruction, we obtained a 2.8 Å resolution structure and confirmed that the sheath is composed of Cis2 (Figure 1C/D). Except for the flexible region between domains 2 and 3 (residues 209-291), all residues could be built into the final atomic model. We compared the structure of Cis2 from *Hb* with homologs from *Serratia entomophila* (*21*), *Algoriphagus machipongonesis* (*24*) and *Photorhabdus asymbiotica* (*22*). Domains 1 and 2, which are important for the contractile mechanism, are particularly well conserved across all three representatives, while the surface-exposed domain 3 showed the highest structural variability (Figure S1C/D/E).

### H. borinquense exhibits antagonism towards Haloferax volcanii

To obtain functional insights into the *Hb* CIS, we first generated antibodies against the putative inner tube (Cis1) and sheath (Cis2). Using Western blotting, we then monitored changes in expression over the course of 72 h. Sheath expression remained relatively stable during exponential phase, while protein levels for the inner tube started dropping rapidly after mid-exponential phase (Figure S1F/G). A similar change in abundance of the inner tube was previously observed for the *V. cholerae* T6SS (*28*).

Having identified conditions for CIS expression, we then set out to probe whether *Hb* exerted an antagonistic effect on other organisms using a killing assay. In order to perform killing assays in media that is compatible with both *Hb* and the prey, we chose to use the haloarchaeon *Haloferax volcanii* (hereafter *H. volcanii* or *Hv*). Since *Hv* is genetically tractable and does not harbor a CIS gene cluster, it is an excellent strain for analyzing cell-cell interactions in liquid and on solid media.

To perform killing assays, both strains were first grown to early exponential phase and then mixed together at a ratio of 1:5 (*Hv*:*Hb*). During liquid killing assays, the mixture was incubated shaking for 24 h, and for solid killing assays the mixture was spotted onto solid medium and incubated for 24 h. To quantify these killing assays, cells from liquid and solid killing assays were serially diluted and spotted onto solid medium, which specifically selected for *Hv*. The results of the liquid killing assay showed only a slight decrease of *Hv* growth compared to the control. The solid killing assay, on the other hand, revealed a significant drop in *Hv* growth by more than 1,000-fold (Figure 1E).

### Antagonism is dependent on cell-cell contact and CISs

To better understand this antagonistic behavior, we studied cell-cell interactions on a single-cell level. We applied time lapse imaging to a 1:2 mixture of *Hv* cells expressing cytoplasmic mScarlet (hereafter *Hv*-mScarlet), and *Hb* cells growing at 45°C for 960 min. During cell fate quantification, mScarlet was initially used to identify *Hv-*mScarlet cells at the start of a time series. However, *Hv* mScarlet signal is lost after the first few frames during imaging. We postulate that this occurs due to the presence of *Hb*.

During time lapse imaging, we identified three distinct *Hv*-mScarlet cell fates: “cell lysis”, “no proliferation” and “normal proliferation” (Figure 2A/B/C; Movie S1; Movie S2). We first quantified the fates of *Hv*-mScarlet cells that were in contact with *Hb*. This revealed a significant number of interactions, in which *Hv*-mScarlet cells lysed (41.6%, n = 332) or no longer proliferated (43.7%, n = 332), with a total 85.3% of the interactions being antagonistic (Figure 2D). In contrast, *Hv*-mScarlet cells that were not in contact with *Hb* during imaging primarily underwent normal cell proliferation (95.1%, n = 243), with only 4.9% of the cells lysing or not proliferating. This suggests that the observed antagonistic effect on *Hv*-mScarlet relies on direct cell-cell contact with *Hb*, which is consistent with findings from the solid killing assay (Figure 1E).

**Figure 2:**
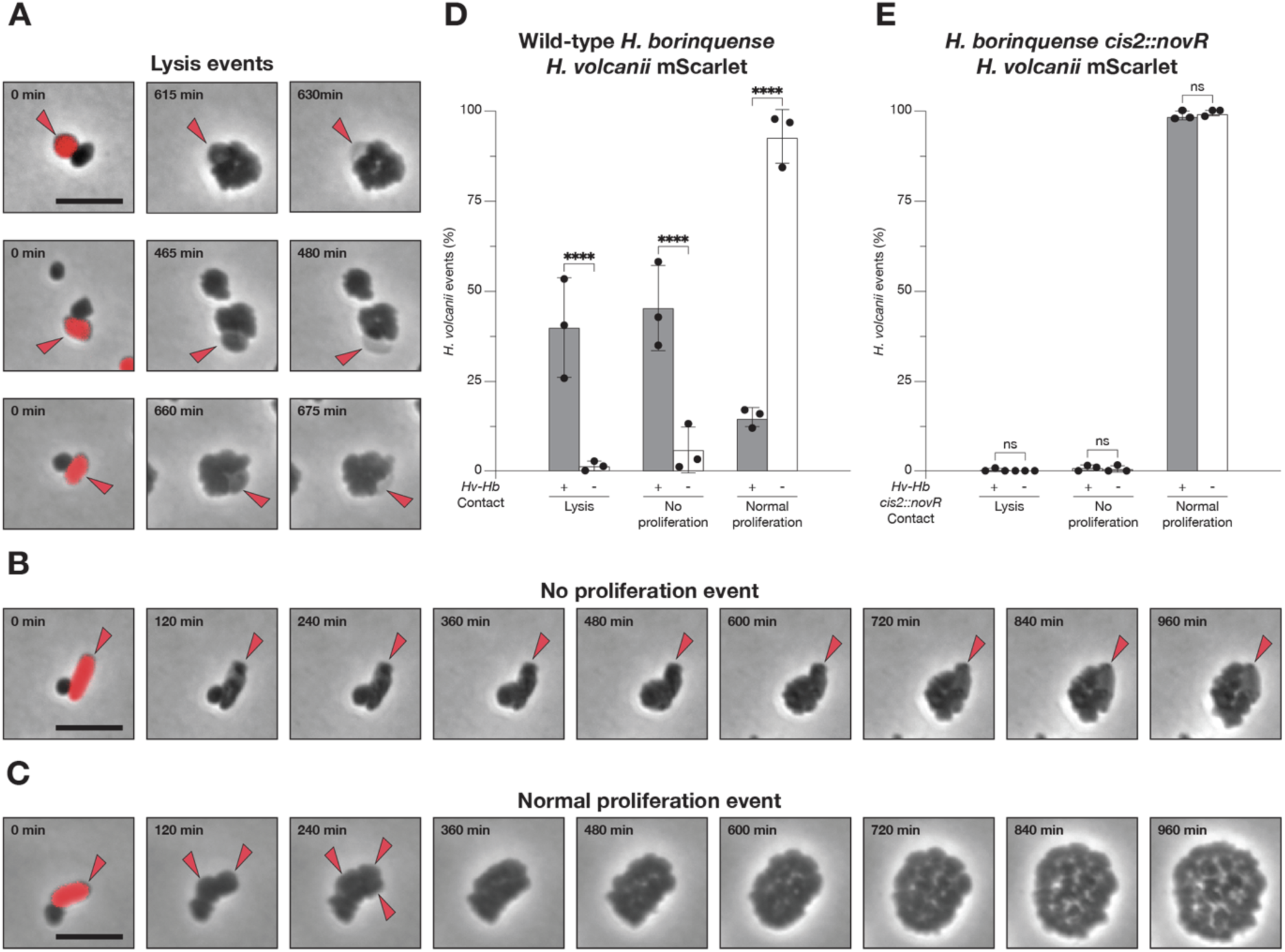
Antagonism is contact and CIS-dependent. **(A-C)** Time lapse imaging (15 min frame rate) of a co-incubation of *H. borinquense* with *H. volcanii* mScarlet during 960 min at 45°C. Shown are selected frames as overlays of phase contrast images with red fluorescence (respective time points are indicated). Red fluorescence originated from mScarlet expressed in *H. volcanii*. Due to the significant drop in mScarlet signal after the first time point, red arrowheads are used to follow the fate of *H. volcanii* mScarlet cells. Three distinct classes of *H. volcanii* cell fates were observed upon cell-cell interactions: Panel **A** shows three examples of “cell lysis” events, characterized by the loss of cellular integrity. Panel **B** shows a “no proliferation” event, in which wild-type *H. borinquense* is observed to proliferate, while *H. volcanii* mScarlet proliferation is impaired. Panel **C** shows a “normal proliferation” event, in which both strains are observed dividing during imaging. The growth of *Hv* was only followed with red arrowheads in the first three images. Note that for panel **C** the experiment was done with the sheath mutant *H. borinquense* c*is2*::*novR*. Scale bar: 5 µm. **(D)** Quantification of the three cell fates exhibited by *H. volcanii* mScarlet with and without direct contact to wild-type *H. borinquense*. Contact events between the two strains saw a significant decrease in the number of “normal proliferation” events with a significant increase in both “cell lysis” and “no proliferation” events (****P < 0.0001; N^contact^ = 243; N^no^ ^contact^ = 332; mean with SD). 2way ANOVA analysis was performed to determine the statistical significance of contact in perturbing *H. volcanii*. **(E)** Quantification of the three cell fates exhibited by *H. volcanii* mScarlet with and without direct contact to *H. borinquense* c*is2*::*novR*. Contact between the two strains did not have an influence on the number of *H. volcanii* mScarlet cells with “normal proliferation” events and did not lead to a significant increase in the number of “cell lysis” nor “no proliferation” events (ns > 0.9999; N^contact^ = 343; N^no^ ^contact^ = 320; mean with SD). 2way ANOVA analysis was performed to determine the statistical significance of contact in perturbing *H. volcanii*.

To determine the role of the CIS in this antagonistic interaction, we set out to disrupt the CIS gene cluster and test the resulting mutant in time lapse imaging. While there are no established genetic tools for manipulating *Hb*, closely related haloarchaea are known to perform homologous recombination (*29*). Therefore, by transforming a plasmid with a novobiocin resistance gene flanked by sheath gene fragments, we were able to trigger homologous recombination and disrupt the sheath gene in *Hb* (hereafter *Hb-cis2*::*novR*). The absence of proper CIS assembly in the generated mutant was verified via sequencing, sheath purifications, where the typical sheath structures were no longer observable, and Western blots that showed that the full-length sheath protein was no longer expressed (Figure S2). In addition, Western blotting also found that inner tube (Cis1) expression was impaired by the homologous recombination event, likely due to the genes location directly downstream of *cis2*. Together, these findings indicate that the mutant was no longer able to assemble functional CISs. We then tested the interaction between *Hb*-*cis2*::*novR* and *Hv*-mScarlet using time lapse imaging. Strikingly, *Hv*-mScarlet “cell lysis” and “no proliferation” events were almost entirely absent (1.2% for *Hv*-mScarlet cells in contact with *Hb*-c*is2*::*novR*, n = 343; 0.6% for *Hv*-mScarlet cells without contact with *Hb*-c*is2*::*novR*, n = 320) (Figure 2E). These findings suggest that both direct cell-cell contact as well as CIS assembly are required for the antagonistic activity exerted by *Hb* on *Hv-*mScarlet.

### The CIS binds to the archaeal cytoplasmic membrane in a T6SS-like manner

Time lapse imaging highlighted the importance of the *Hb* CIS for contact-dependent antagonism. To understand how the CIS architecture and mode-of-action contribute to its function, a combination of cryo-focused ion beam (cryoFIB) milling and cryo-electron tomography (cryoET) was applied to whole *Hb* cells. Within the resulting tomograms, CISs were observed in both extended (∼16 nm diameter) and contracted (∼19 nm diameter) states (Figure S3A/B). Extended CIS structures, in which the baseplate was identifiable, were always perpendicular to the cytoplasmic membrane with a putative membrane anchoring structure (Figure 3A/C). Contracted sheaths were observed as either free floating or perpendicular to the cytoplasmic membrane (Figure 3B). Based on the T6SS-like sub-cellular localization of the CIS, hereafter we will refer to the system as an archaeal T6SS.

**Figure 3:**
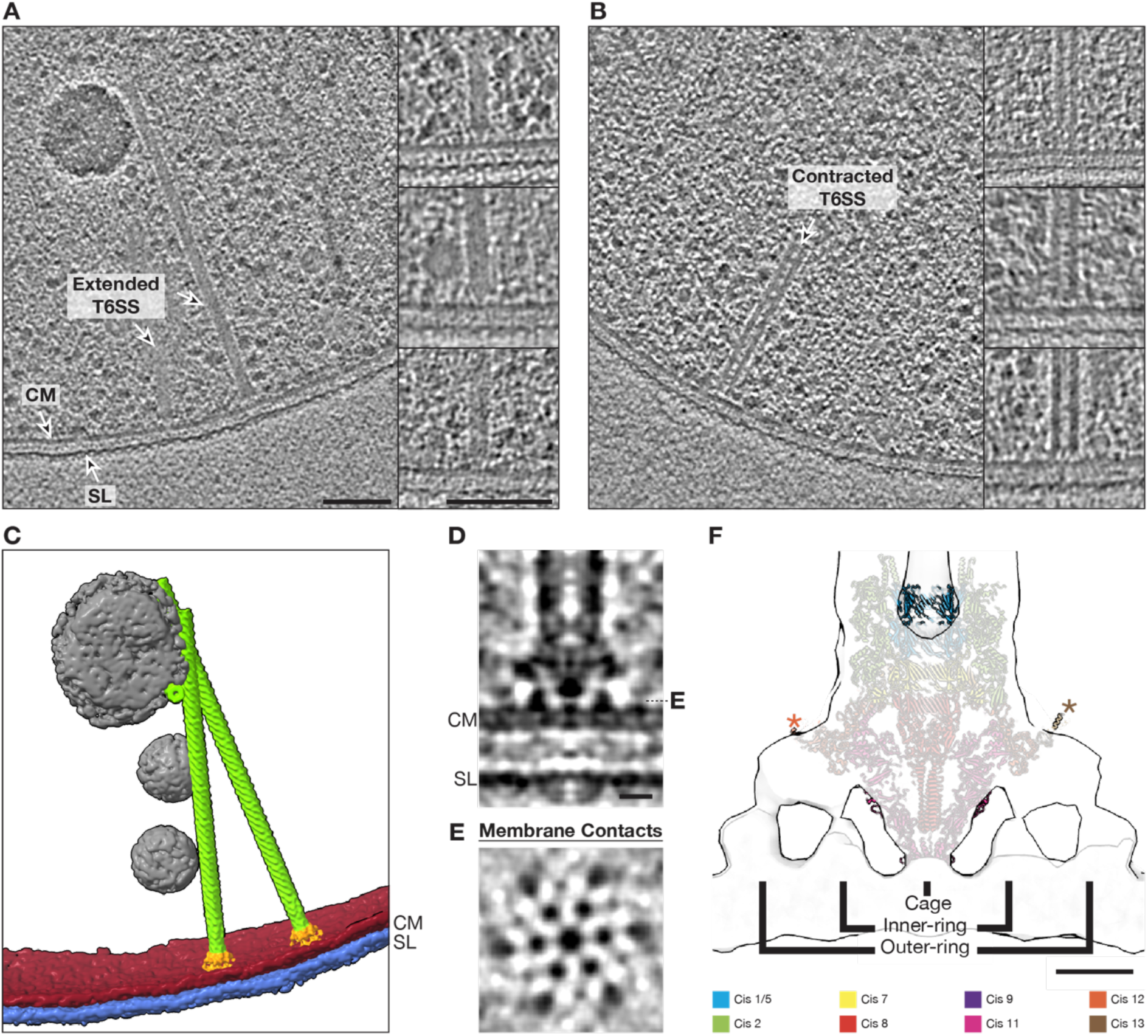
CIS binds to archaeal membrane in a T6SS-like manner. **(A/B)** Slices through tomograms collected on cryoFIB-milled *H. borinquense* cells grown at 45°C and 2.45 M NaCl. Tomograms reveal extended (panel **A**) and contracted (panel **B**) T6SS-like CIS structures inside the cells. Additional examples of both extended and contracted T6SSs are shown on the right. Shown are 13.6 nm thick slices through Tegunov deconvolution-filtered tomograms. CM: cytoplasmic membrane, SL: S-layer. Scale bar: 100 nm. **(C)** Segmentation of the cryo-tomogram depicted in panel **A**, with the cytoplasmic membrane (SM) in red, S-layer (SL) in blue, putative storage granules in grey, and the T6SS baseplate and sheath-tube module in orange and green, respectively. **(D/E)** Subtomogram average of C6-symmetrized extended T6SS baseplate and membrane anchor using 57 particles that were extracted from filtered tomograms. Shown are 1.36 nm thick longitudinal (panel **D**) and perpendicular slices (panel **E**). The perpendicular slice reveals contacts with the membrane, i.e. densities of six ‘feet-like’ structures that are arranged around the central ‘cage’. CM: cytoplasmic membrane, SL: S-layer. Scale bar: 10 nm. **(F)** Shown is a longitudinal slice through a Gaussian-filtered subtomogram average isosurface (grey) from panels **D/E.** The extended baseplate of the tCIS from *Anabaena* sp. PCC 7120 (PDB: 7B5H, (*27*)) was fitted into the average. tCIS proteins Cis19 and Cis6 were removed since the *H.* borinquense CIS gene cluster lacks homologues of these proteins. Note the good fit of Cis11, which makes contact with the membrane. Cis12 and Cis13 (highlighted with asterisks) at the periphery of the baseplate may be involved in interactions with putative membrane-anchoring proteins. Scale bar: 10 nm.

The archaeal cell envelope of *Hb* is fundamentally different from the architecture in Gram-negative bacteria and is comprised of a cytoplasmic membrane, which is associated with an ordered layer of glycosylated protein – called the S-layer (*30*, *31*). We were therefore interested in how a T6SS evolved to make contact with this distinct type of cell envelope. Unfortunately, the high salt concentrations that are required to propagate *Hb* resulted in a poor signal-to-noise ratio. This made characterizing the structural modules of the T6SS particularly difficult. We set out to improve the contrast of the baseplate region by subtomogram averaging. Due to the structural heterogeneity of contracted T6SSs, possibly due to the presence of disassembly intermediates, we focused on generating an average of the extended state. Initial averages identified a 6-fold rotational symmetry in the sheath-tube, baseplate, and membrane anchoring modules (Figure S3C/D/E). We therefore applied C6 symmetry during the final rounds of subtomogram averaging (Figure S3F). An average of the sheath-tube module, in the extended state, revealed densities that likely represent the sheath and the inner tube with a hollow lumen (Figure S3B).

Resolving the contact between the membrane and the baseplate was still difficult, therefore, subtomogram averaging was performed using deconvolution-filtered data to improve the contrast. The resulting average (Figure 3D/E) showed the same features and allowed better visualization of the baseplate and membrane anchor. Both averages did not allow individual subunits of the baseplate to be resolved, however, six ‘feet-like’ structures emanating from the periphery of the baseplate and a central ‘cage’ at the base of the baseplate are clearly visible. Each individual ‘foot-like’ structure makes contact with the archaeal cytoplasmic membrane in two locations, forming an outer and inner ring of membrane attachment points, which center around the ‘cage’. In addition, the cage also forms contact with the cytoplasmic membrane at the center of the T6SS.

To identify the proteins likely involved in these membrane contact points, the high-resolution structure of the pre-firing baseplate from *Anabaena* sp. PCC 7120 (hereafter tCIS) (*27*) was docked into the subtomogram average from deconvolved data (Figure 3F). The tCIS was chosen due to the presence of all the structural proteins found in the *Hb* CIS gene cluster, and because it also anchors itself to membranes *in situ*. In addition, it is one of the most closely related CIS gene clusters that has been studied.

The structures showed a high overall structural similarity at the baseplate (Cis1/2/5/7/8/9/12/13) and the ‘cage’ (Cis11), and allowed the atomic model to be docked into the subtomogram average. The fitting also permitted us to postulate that Cis12 and Cis13 are likely involved in membrane attachment, as both fit within the peripheral densities of the baseplate. An additional contact made at the base of the baseplate correlates well with the ‘cage’ formed by Cis11. Due to the presence of two *cis11* genes in the *Hb* gene cluster, the cage is likely formed by either one or both of these gene products. From the subtomogram average it was impossible to determine how the CIS attaches to the membrane. Cis11, 12 and 13 may interact directly with the membrane or via a membrane-embedded protein that was not resolved during averaging.

This mechanism of attaching to membranes via six ‘feet-like’ structures is also observed for the T6SS subtype 4 (T6SS*^iv^*) found in *Candidatus* Amoebophilus asiaticus (*32*), however, the overall architecture of membrane anchoring looks drastically different.

### The T6SS gene cluster encodes putative toxin-immunity pairs

In many cases, T6SSs harbor toxins that are believed to be translocated upon contraction (*33*). Analysis of the gene cluster identified an ‘interspacing’ region in the middle of the gene cluster which contains nine putative genes. To analyze if these proteins had potential toxic activity, we expressed each gene in *Hv* using a plasmid for constitutive expression and C-terminal His-tagging. Transformation of the generated plasmids into *Hv* was successful for all genes, except for *Hbor_38740* and *Hbor_38760* (Figure S4A). Repeated attempts to transform plasmids with these two genes either resulted in the absence of any transformants, or in transformants that contained a plasmid where the respective gene was excised. Bioinformatic analysis of both genes were only able to identify one putative domain at the N-terminus of *Hbor_38740*. This DUF4157 domain has been previously identified to be associated with CIS-related cargo effectors (*24*, *34*).

Interestingly, many T6SS toxin genes are accompanied by an immunity gene in order to prevent self-intoxication and toxicity from sister cells (*35*). Within the interspacing region, *Hbor_38740* and *Hbor_38760* are closely followed by two downstream genes of unknown function. When we designed constructs to express *Hbor_38740* and *Hbor_38760* in tandem with their downstream partners *Hbor_38750* and *Hbor_38770*, respectively, we found that the constructs were successfully transformed and expressed (Figure S4B). Altogether, our data indicates that the *Hb* T6SS interspacing region encodes two putative toxin-immunity pairs.

### Other haloarchaeal species express CIS and exhibit contact-dependent antagonism

The characterization of an archaeal T6SS in *Hb* revealed a new type of mechanism by which archaea can interact with their environment. To determine whether the observed function and mode-of-action are conserved, we explored the abundance of CIS gene clusters in archaea. We focused on identifying CISs in the class *Halobacteria* (halophilic archaea), for which we have established functional assays to characterize archaea-archaea interactions. From a total of 350 analyzed genomes, 41 strains (12 %) harbor a gene cluster that contains the structural genes required for assembling a functional CIS (*cis*1-16) (Figure 4A). These gene clusters were detected in three out of four phylogenetic orders (Figure S5A), with all representatives having a rather well conserved gene content and synteny (Figure 4A). The phylogenetic trees generated from 16S rDNA and Cis11a sequences did not match (Figure 4A; Figure S5A), indicating horizontal gene transfer of CIS gene clusters among halophilic archaea. We could verify that sheath-like structures were expressed in five of the identified strains (bold species names in Figure 4A).

**Figure 4:**
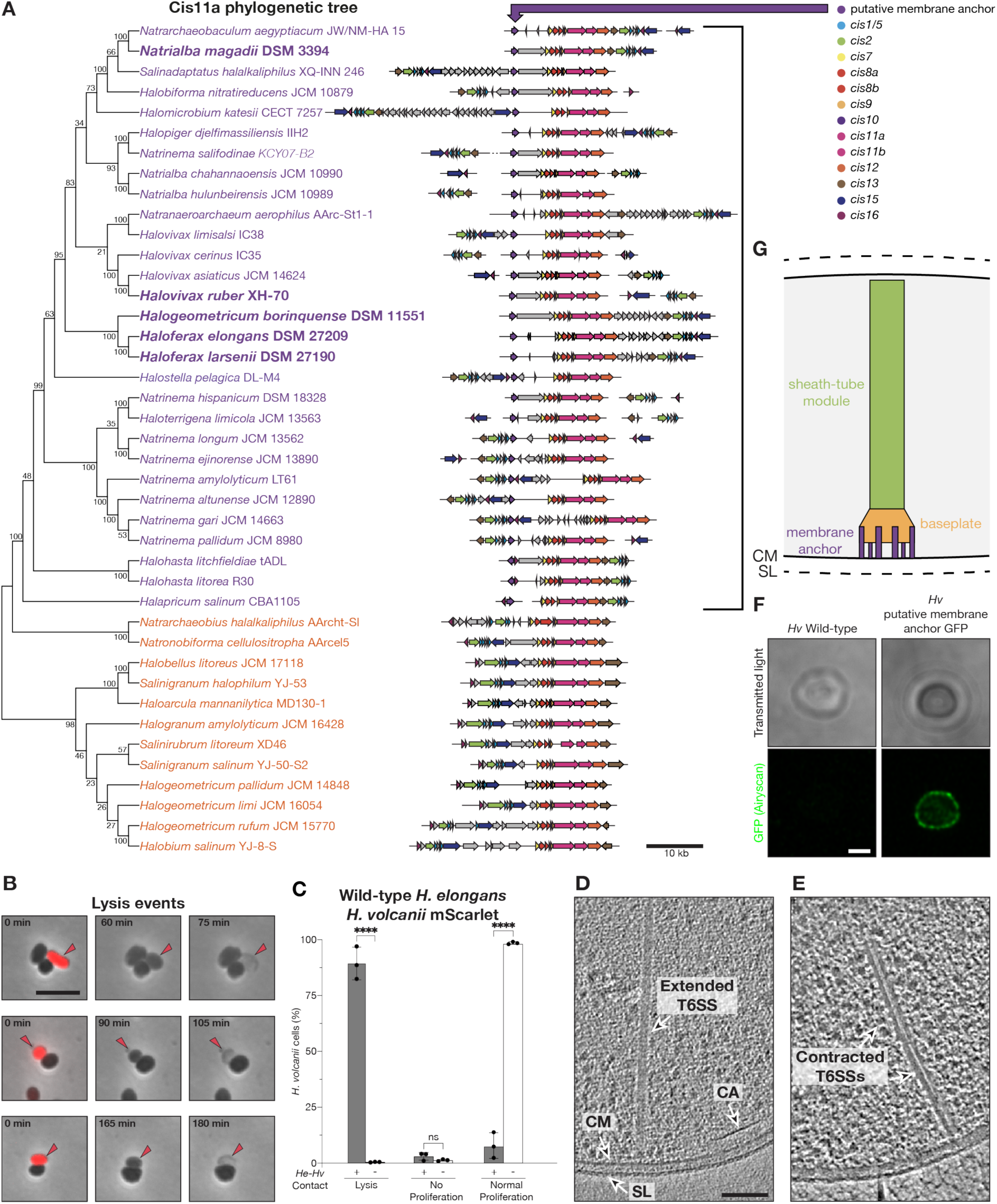
CIS gene clusters are conserved in haloarchaea with a subset functioning via a contact-dependent T6SS mode-of-action. **(A)** Phylogenetic tree (bootstrap values shown at nodes) based on Cis11a homologs from haloarchaea paired with schematics of the corresponding CIS gene clusters. The tree reveals two stable clades (indicated in purple and orange). The gene clusters have a general synteny and an additional conserved putative membrane protein in CISs from the purple clade. Species, for which the expression of CIS-like structures was verified by negative stain EM, are shown in bold font. **(B)** Time lapse imaging (15 min frame rate) of a co-incubation of *H. elongans* and *H. volcanii* mScarlet during 960 min at 45°C. Shown are selected frames as overlays of phase contrast images with red fluorescence (respective time points are indicated). Red fluorescence originated from mScarlet expressed in *H. volcanii*. Due to the significant drop in fluorescence signal after the first time point, red arrows are used to follow the fate of *H. volcanii* mScarlet cells. The panel shows three examples of “cell lysis” events, characterized by the loss of cellular integrity. Scale bar: 5 µm. **(C)** Quantification of the three cell fates (“cell lysis”, “no proliferation” and “normal proliferation”) exhibited by *H. volcanii* mScarlet with and without direct contact to *H. elongans*. Contact events between the two strains saw a significant increase in “cell lysis” events (****P < 0.0001; N^contact^ = 321; N^no^ ^contact^ = 1235; mean with SD). 2way ANOVA analysis was performed to determine the statistical significance of contact in perturbing *H. volcanii* mScarlet. **(D/E)** Slices through tomograms collected on cryoFIB-milled *H. elongans* cells grown at 45°C and 2.45 M NaCl. Panel **D** shows a T6SS-like CIS structure in the extended state. Extended T6SSs are identifiable in the cytoplasm and attached to the cytoplasmic membrane with an architecture similar to that seen in *H. borinquense*. Panel **E** shows the T6SS in the contracted state. Contracted T6SS structures are found both free floating or perpendicular to the cytoplasmic membrane, similar to *H. borinquense*. Shown are 13.6 nm cross-sections through Tegunov deconvolution-filtered tomograms. CM: cytoplasmic membrane, SL: S-layer, CA: chemotaxis array. Scale bar: 100 nm. **(F)** Representative confocal microscopy images of wild-type *H. volcanii* and *H. volcanii* expressing the putative membrane anchor *Hbor_38620* tagged with GFP. The top row of images show the overall cell shapes via a minimum intensity *z*-projection of the transmitted light channel. The bottom row of images show a single slice of the fluorescent GFP signal from the same cells. GFP signal was not observed in the wild-type strain, however, the putative membrane anchor is observed to localize to the cell membrane. Scale bar: 1 µm. **(G)** Schematic of the haloarchaeal T6SS. CryoET imaging, functional assays, Cis11a phylogenetics and gene cluster analyses lead us to hypothesize that representatives of the purple clade may all have a T6SSs mode-of-action. We postulate that this mode-of-action is facilitated by membrane anchoring via the identified conserved putative membrane-anchoring protein (homologs of *Hbor_38620*).

Among the identified CIS-positive strains, *Haloferax elongans* (hereafter *H. elongans* or *He*) showed the highest abundance of observable sheath structures. Since *He* grows under the same conditions as *Hv*, we proceeded to study their interaction by time lapse imaging. Similar to our observations of co-incubating *Hb* with *Hv-*mScarlet (Figure 2), a co-incubation of *He* with *Hv-* mScarlet (2:1 ratio) revealed “cell lysis”, “no proliferation” and “normal proliferation” cell fates (Figure 4B/C). Interestingly, contact with *He* led to a significantly higher percentage of *Hv*-mScarlet “cell lysis” events (89.1%, n = 321 for *He* vs 41.6%, n = 332 for *Hb*), with a total of 92.2% of interactions being antagonistic (Figure 4C). *Hv-*mScarlet cells growing without coming in contact with *He* had an insignificant number of antagonistic-like cell fates (1.3%, n = 1235). It should be noted that the conditions during imaging were not optimal for *He*, and led to occasional cell lysis of *He* even in the absence of *Hv*-mScarlet (Figure S6).

The contact-dependent antagonism indicated the possibility of a T6SS mode-of-action in *He*. We verified this theory by imaging *He* cells by cryoET. The resulting tomograms showed *He* T6SSs in both extended and contracted states. For structures in the extended state, where the baseplate was visible, the T6SSs were always attached to the cytoplasmic membrane in a visually similar manner as observed for the *Hb* T6SS (Figure 4D). T6SSs in the contracted state were less common (Figure 4E). Together, the time lapse imaging and *in situ* structural data indicate that the *He* CIS is an additional example of an archaeal T6SS that likely mediates contact-dependent antagonism.

### The T6SS mode-of-action correlates with a conserved putative membrane anchor

The identification of two CIS gene clusters with a T6SS mode-of-action indicates an evolutionary drive towards anchoring these systems to the archaeal cell envelope. Therefore, we further analyzed the haloarchaeal CIS gene clusters for potential membrane-anchoring proteins. The Cis11a phylogenetic tree revealed two distinct, stable groups of representatives (labeled purple and orange, respectively in Figure 4A). Interestingly, the gene clusters corresponding to the first group of 29 representatives (including Cis11a from *Hb* and *He*) were always associated with an uncharacterized conserved gene. In *Hb* and *He*, this gene (*Hbor_38620* and *SAMN04488691_10831*, respectively) is found upstream of the core CIS gene cluster. Bioinformatic characterization of *Hbor_38620* identified six predicted transmembrane helices and a ∼150 residue-long cytoplasmic domain (Figure S5B/C), indicating a putative role in anchoring the T6SS to the cytoplasmic membrane. The absence of *Hbor_38620* homologs in the second group of CIS gene clusters (orange in Figure 4A) may suggest a different mode-of-action for these systems.

To test whether *Hbor_38620* functions as a membrane protein, we GFP-tagged and expressed *Hbor_38620* in *Hv*. Using confocal microscopy, we observed that *Hbor_38620* localized to the cell membrane (Figure 4F). The localization of *Hbor_38620* at the cell membrane, coupled with the association of *Hbor_38620* homologues to CIS gene clusters with a T6SS mode-of-action could hint a potential role in anchoring the archaeal T6SS to the membrane (Figure 4G).

## Discussion

Archaea are known to interact with their environment, however, the mechanisms that govern these interactions are largely unclear. The identification of bacteria-like CIS gene clusters within the archaeal domain allows new avenues for understanding how archaea interact with their environment to be explored. In this work, we studied an archaeal CIS in *Halogeometricum borinquense*. Characterization of both its structure and function gives new insights into the evolution of T6SSs and furthers our understanding of the roles that archaea have in their environment.

Our first major insight was that the CIS gene cluster in *Hb* has a T6SS mode-of-action, i.e. it functions while localized in the cytoplasm and bound to the cytoplasmic membrane. However, its architecture is different to the previously described T6SSs. In the canonical T6SSs (subtype i-iii; (*36–38*) and the recently reported T6SS subtype iv found in *Aureispira* sp. CCB-QB1 (*39*), a large trans-envelope complex spans between the inner and outer membrane. Likely as an adaptation to the archaeal cell envelope, which features an S-layer instead of a cell wall and outer membrane, the T6SS in *Hb* does not seem to have a complex outside of the cytoplasmic membrane. The system therefore appears rather minimal, akin to the T6SS subtype iv in the Gram-negative bacteria *Candidatus* Amoebophilus asiaticus (*32*). Despite this overarching similarity, however, the interactions between baseplate and membrane are different. Subtomogram averaging of the *Hb* T6SS revealed that the interactions between the T6SS and the membrane are mediated via the ‘cage’, which surrounds the spike, as well as by six ‘feet-like’ densities emanating from the periphery of the baseplate (Figure 3E). In the T6SS*^iv^* of *Amoebophilus,* the ‘feet-like’ structures emanate from the base of the baseplate rather than from the periphery as seen in *Hb*.

From the *Hb* T6SS subtomogram average, we also identified that the feet-like densities are likely not fully accounted for by any of the conserved baseplate components (Figure 3F). We speculate that the conserved putative membrane anchor (*Hbor_38620*) may be involved in these interactions. From phylogenetic analysis of 41 haloarchaeal CIS gene clusters, we identified a distinct subset of 29 strains that all share this putative membrane protein, including *He*, which we experimentally showed to also acts as a T6SS. Interestingly, homologs of this putative membrane anchor are only identifiable in haloarchaea with CIS gene clusters. Our data therefore indicates that this membrane protein may be useful for predicting a T6SS mode-of-action in uncharacterized CIS gene clusters of haloarchaea.

Our second major finding was that *Hb* disrupts proliferation and induces “cell lysis” of *Hv* in a contact-and T6SS-dependent manner. Interestingly, this contact-dependent antagonistic behavior was also verifiable in *He*. Antagonism has been observed in other archaeal species, for example, through the secretion of small peptide archaeocins into their surrounding environment (*9*, *11*). However, the targeted delivery of toxins from one archaeon to another has not been observed to date. Such a toxin delivery system has implications in the study of archaeal biofilms and other niches, in which archaea reside in close contact (*4*).

During the analysis of the T6SS gene cluster, we also identified two toxin-immunity pairs in the interspacing region. The presence of toxin-immunity pairs has only been identified in canonical T6SSs (*20*, *40*). From tomography data and subtomogram averaging, we did not observe extra densities within the lumen of the inner tube. We therefore postulate that these toxins attach to the T6SS spike, as was previously shown (*41*, *42*).

In conclusion, the insights and methods used in this study provide a framework for studying novel types of archaeal cell-cell interaction in the future. It will be exciting to explore the ecological advantage that haloarchaea may have acquired by expressing CIS gene clusters in complex microbial communities. In particular, the identification of CIS gene clusters in more distant relatives, including methanogenic and anaerobic methanotrophic archaea (*11–13*), which thrive in close interactions with other species, indicates, that these systems could provide a competitive advantage within a plethora of different ecological niches.

## Materials and Methods

### Strains and culturing conditions

The archaeal strains studied in this work and their growth conditions are found in Table S1. The archaeal media YPC and Ca are based on the previously described *Hv*-YPC and *Hv*-Ca media, respectively (*43*). Growth media with a DSM number are from the German collection of microorganisms and cell cultures. Cultures were grown in 5 ml aeration tubes and larger volumes were grown in flasks at 200 rpm shaking unless stated otherwise. Listed growth media were solidified, when necessary, with 1.5-2% agar. Archaeal strains used in functional assays and the generated *Hb-*c*is2*::*novR* strain were selected for with varying concentrations of the antibiotics lovastatin (LovR) and novobiocin (NovR). Additional information about the growth of each strain can be found in the preceding methods sections. The *E. coli* strain Top10 was used to generate and propagate all constructs used in this study. Dam and Dcm methylase deficient K12 *E. coli* was used to prepare unmethylated DNA required for archaeal transformations. Both bacterial strains were grown in LB media or agar substituted with 100 µg ml^−1^ ampicillin.

### Plasmids and genetic manipulation of haloarchaea

All plasmids that were generated in this work are based on the backbones of plasmid pWL102 (DSM 5717) or pET15b (Novagen). The full list of generated plasmids and the oligonucleotides used in this work are summarized in Table S2 and S3, respectively. Transformations of plasmids into *Hv* and *Hb* were performed as previously described (*43*). In brief, cells were spheroplasted in a spheroplasting solution (1 M NaCl, 27 mM KCl, 50 mM Tris HCl pH 8.5, 15% sucrose) with 50 mM EDTA. Subsequently, 30 µl of DNA solution (10 µl ∼1-2 µg dam^−^ DNA, 15 µl unbuffered spheroplasting solution [1 M NaCl, 27 mM KCl, 15% sucrose, pH 7.5], 5 µl 0.5 M EDTA pH 8.0) was mixed with the spheroplasted cells. To begin the transformation, equal volume PEG600 (60% PEG600, 40% unbuffered spheroplasting solution) was gently added to the cells. Spheroplasting was stopped by adding a spheroplasting dilution solution (3.1 M NaCl, 113.2 mM MgCl_2_·6H_2_O, 108.9 mM MgSO_4_·7H_2_O, 72 mM KCl, 15.3 mM Tris.HCl pH 7.5, 15% sucrose, 3 mM CaCl_2_). Cells were then added to regeneration solution (2.5 M NaCl, 88.6 mM MgCl_2_·6H_2_O, 85.2 mM MgSO_4_·7H_2_O, 56.3 mM KCl, 12 mM Tris.HCl pH 7.5, 1.25 g yeast extract, 0.25 g peptone, 0.25 g casamino acids, 15% sucrose, 3 mM CaCl_2_). To recover, cells were left to incubate at 45°C for 2 h without shaking and subsequently 4 h shaking. YPC solid media supplemented with lovastatin or novobiocin was used to select for successful transformations. Transformation of plasmids, with a pWL102 backbone, used selective media containing either 4 µg ml^−1^ or 0.5 µg ml^−1^ lovastatin. Plasmids with a novobiocin resistance gene were selected for with 0.3 µg ml^−1^ novobiocin. To verify that plasmids had been successfully transformed or that homologous recombination occurred PCR was used.

### Sheath preparation of haloarchaea for negative stain and mass spectrometry

To isolate sheath, archaeal strains were grown for 24 h (*Hb* and *Hb-*c*is2*::*novR)* or 72 h (CIS expression tests) in 25 ml of their respective media (Table S1). Cells were then pelleted (7,000 *g*, 10 min) and resuspended in 3-5 ml lysis buffer (150 mM NaCl, 50 mM Tris-HCl, 0.5xCellLytic B [Sigma-Aldrich], 1 % Triton X-100, 50 µg ml^−1^ DNAse I, pH 7.4) and incubated for 30-60 min at 37°C. Cell debris was removed via centrifugation at 15,000 *g* for 15 min at 4°C. To isolate the sheath structures, the supernatant was ultracentrifuged at 150,000 *g* for 1 h at 4°C. The resulting pellet was resuspended in 25-50 µl of resuspension buffer (150 mM NaCl, 50mM Tris, pH 7.4) and immediately used for electron microscopy. To identify proteins in the sheath preparation, samples were sent for mass spectrometry analysis at the Functional Genomics Center Zürich. Samples were prepared for mass spectrometry by precipitating proteins in a 30 µl sample using 10% trichloroacetic acid. The resulting precipitant was pelleted and washed twice with cold acetone. After drying the pellet, it was dissolved in 45 µl buffer (10 mM Tris, 2 mM CaCl_2_, pH 8.2) and 5 µl trypsin (100 ng/µl in 10 mM HCl) to digest the protein and then microwaved for 30 min at 60°C. Samples were dried and dissolved in 20 µl 0.1% formic acid, diluted 1:10 and transferred to autosampler vials. 2µl were injected for liquid chromatography with tandem mass spectrometry analysis. Database searches were performed by using Mascot (swissprot, all species and trembl_archaea) search program. For search results, stringent settings were applied in Scaffold (1% protein false discovery rate, a minimum of 2 peptides per protein, 0.1% peptide false discovery rate). Mass spectrometry results were visualized with the software Scaffold (Proteome Software, v5.0.1).

### Negative stain electron microscopy

3 µl of sheath preparation sample was incubated for 1 min on glow-discharged carbon-coated formvar grids (Electron Microscopy Sciences). To stain the sample, 3 µl of 1% phosphotungstic acid was added to the grid three times and left to incubate 0, 15 and 30 s. To image the grids, an 80 kV Thermo Fisher Scientific Morgagni transmission electron microscope was used.

### Single particle cryoEM of the contracted H. borinquense sheath

Samples for single particle cryoEM analysis were prepared from a crude sheath preparation of *Hb* in mid-exponential phase. The purified samples were directly used for vitrification with a Vitrobot Mark IV (Thermo Fisher Scientific) at 8°C and 100% humidity. 4 µl sample was applied onto Au R2/2 200 mesh (Quantifoil) grids, and excess liquid was then blotted away twice (2.5 s) after waiting for 15 s. The sample was then immediately plunged into liquid ethane/propane (37% v/v ethane). The vitrified grids were used for single particle data collection.

Data collection was performed on a Titan Krios (Thermo Fisher Scientific) transmission electron microscope equipped with a K3 direct electron detector (Gatan) and Quantum LS GIF (Gatan, 20 eV slit). Data collection was performed using SerialEM (Mastronarde, 2005). The potential contracted sheath particles were manually picked in low magnification and were used as reference to align and collect data at high magnification (nominal magnification 81,000 x). Data was collected as a movie stack with 50 frames at super-resolution mode (an effective pixel size of 0.55 Å). The exposure time was 2.5 s, with an accumulated dose of ∼65 e^−^/Å^2^. The movie frames of each collected stack were aligned and summed up into one micrograph with dose weighting at a binning factor of 2 using MotionCorr2 (pixel size of 1.10 Å) (*44*). The CTF parameters were then estimated using Gctf (*45*). A total of 638 micrographs, with a defocus value ranging from −1 µm to −3.3 µm were used during image processing.

The image processing was performed similar to previous reports (*24*, *27*). Briefly, the contracted sheaths were manually picked using Relion3.0 (*46*) with start–end coordinate pairs. The filaments were segmented with the inter-box distance of 18.3 Å. Bad particles were removed through one round of 2D classification at a binning factor of 4, and particles in good 2D classes were used for 3D auto-refinement applying 6-fold symmetry but not helical symmetry. The initial helical parameters were then interpreted in real space using the *relion_helix_toolbox* and were further optimized in the following helical reconstruction (*47*). A total of 127,891 particles were used to reconstruct the contracted *Hb* sheath structure at a resolution of 3.4 Å, which was estimated based on the gold-standard Fourier Shell Correlation (FSC) = 0.143 criteria (*48*). One additional round of CTF parameter optimization was performed and the resolution was improved to 2.9 Å, assuming 6-fold and helical symmetry (twist = 33.24°, rise = 17.09 Å).

### Atomic model building

The quality of the final reconstructed map was shown to be sufficient for *de novo* modelling. The atomic model of the contracted sheath was manually built using COOT (*49*). The model was iteratively refined using RosettaCM (*50*) and *phenix.real_space_refine* (*51*). The final atomic model was validated using *phenix.molprobity* (Table S4). Structural illustrations and superpositions were performed using Chimera (*52*).

### Growth curve measurements

Late exponential phase cultures of *Hb* and *Hb-*c*is2*::*novR* were used to start fresh 25 ml YPC cultures at an OD_600 nm_ of 0.01. Cultures were grown at 45°C, over 72 h. Every 6 h, a sample was taken from the culture to measure the OD_600 nm_. To prepare samples for Western blotting, cells were pelleted to have 50 µl of OD_600 nm_ 1.0. The pellet was resuspended in 1x Laemmli sample buffer (Bio-Rad), denatured for 10 min 95°C and stored at 4°C for Western blotting. Growth curve measurements were plotted using GraphPad Prism (v9.5.1).

### SDS-PAGE and Western Blotting

A standard procedure was followed for running SDS-PAGE and Western blots. In brief, samples were prepared to be at an OD_600 nm_ 1.0 in 50 µl 1x Laemeli buffer (Bio-Rad). Samples were denatured for 10 min at 95°C and then 10 µl were loaded onto a precast 4-15% protein gel (Bio-Rad). Electrophoresis was performed by applying a constant voltage of 200 V in SDS running buffer (Tris/Glycine/SDS, Bio-Rad) for ∼1 h. Samples were then transferred to a nitrocellulose membrane at 100 V for 1 h at 4°C. The membrane was blocked with 5% milk TBST (10 mM Tris-HCl pH 8.0, 150 mM NaCl, 0.05% Tween-20) at room temperature for 1 h. For His-tagged samples, membranes were incubated with monoclonal primary anti-6xHis antibody (Sigma-Aldrich) in 1% milk TBST at 1:3,000. To identify the expression of the *Hb* sheath (anti-Cis2) or inner tube (anti-Cis1) polyclonal antibodies were generated against *Hbor_38820* and *Hbor_38830*, respectively (GenScript). These primary antibodies were incubated in 1% milk TBST at 1:400. After incubation with the primary antibody for 2 h at room temperature, the membranes were washed with TBST 10 min three times. The secondary antibodies, anti-mouse for anti-6xHis (Merk) and anti-rabbit for anti-Cis1/2 (Invitrogen), conjugated with HRP were incubated with the membrane in 1% milk TBST at 1:10,000. After incubating at room temperature for 1 h, the membranes were washed with TBST three times for 10 min. Membranes were imaged (ChemiDoc, Bio-Rad) after applying a chemiluminescent substrate (ECL, Bio-Rad).

### Solid and liquid based killing assay

Late exponential phase cultures were used to start 25 ml YPC media cultures of *Hb* pWL102 and *Hv* pWL102et NovR at OD_600 nm_ 0.01 with lovastatin (0.5 µg ml^−1^). Cultures were grown at 45°C for 24 h. After 24 h, *Hb* and *Hv* OD_600 nm_ was measured and the strains were concentrated to 0.5 and 0.1, respectively. 500 µl of *Hb* culture was then mixed with 500 µl *Hv*. The mixtures were incubated for another 15 min before spotting 100 µl of each mixture onto YPC plates with 0.5 µg ml^−1^ lovastatin. The remaining 900 µl mixtures were left at 45°C shaking for 24 h (liquid killing assay). After the spots dried, the plates were also incubated at 45°C for 24 h (solid killing assay). Each of the liquid killing assay mixtures were serial diluted (10^−2^, 10^−3^, 10^−4^, 10^−5^, 10^−6^) and 25 µl of each dilution was spotted on YPC plates supplemented with 0.3 µg ml^−1^ novobiocin. The spots of the solid killing assay were resuspended with 1 ml of YPC media, serial diluted and spotted on YPC plates supplemented with 0.3 µg ml^−1^ novobiocin in the same manner as the liquid killing assay. After the spots dried the plates were incubated at 45°C for about a week before taking colorimetric images (ChemiDoc, Bio-Rad).

### Time lapse imaging functional assays

Late exponential phase cultures were used to start 25 ml Ca media cultures of *Hb* and *Hv-* mScarlet at OD_600 nm_ 0.01. Cultures were grown at 45°C for 24 h. After 24 h, *Hb* and *Hv* were concentrated to an OD_600 nm_ of 1.0 and 0.5, respectively. Each sample was left to equilibrate at 45°C for 15 min before mixing the two strains together for another 15 min. 7.5 µl of the mixture were spotted onto an agarose pad (2.3 M NaCl, 88.3 mM MgCl_2_·6H_2_O, 85.3 mM MgSO_4_·7H_2_O, 56.5 mM KCl, 12 mM Tris.HCl pH 7.5, 3 mM CaCl_2_ 1% agarose) and left to dry at room temperature. After 30 min, the agarose pad was placed in a glass bottom imaging dish (µ-Dish 35 mm high, Ibidi) and incubated at 45°C for 30 min. To prevent water evaporation and salt crystal formation while imaging, the imaging dish was closed with vacuum grease and parafilm. For imaging the sample, the Thunder Imager 3D cell culture microscope (Leica) i8 CO2 incubator (Pecon) was pre-heated to 45°C for at least 2.5 h before imaging. To identify areas with good coverage of both *Hb* and *Hv-*mScarlet, a tile scan was performed using the Leica Application Suite X (LasX) software (v3.7.6.25997) using the HC PL APO 100x/1.40-0.70 oil objective (Leica). Images were collected using phase contrast to identify all cells and fluorescence (575 nm excitation) to identify mScralet-expressing *H*v-mScarlet. *Hv-* mScarlet signal was quickly lost during imaging and therefore was only used in the first frame to determine which cells were *Hv*. Around 10 target areas were selected for time lapse imaging.

At each area a *z*-stack (2 µm, 9 slices) was collected every 15 min over 16 h. Fiji was used to make maximum intensity projections and to help in the quantification of the time lapse imaging data (*53*). Imaging of the *Hb*-c*is2*::*novR* and *He* followed the same procedure. For the *Hb*-c*is2*::*novR*, however, the initial culture was supplemented with 0.3 µg ml^−1^ novobiocin, however, novobiocin was not present during imaging. Graphs and statistical analyses were performed in GraphPad Prism. To quantify events, antagonism was characterized as an event in which *Hv* no longer proliferated or lysed without producing any daughter cells. Normal proliferation was described as an event where *Hv* continues to proliferate and generate at least one daughter cell.

### Vitrification of samples for cryoET

Samples for tomography data collection were prepared by growing *Hb* or *He* for 6 - 24 h (∼0.1-1 OD_600 nm_) and then concentrating the cells to ∼10-30 OD_600 nm_ to create a lawn of cells on the generated grids. 4 µl of the concentrated cells were then spotted onto a glow discharged copper grid (R2/2, Quantifoil). By using a Vitrobot Mark IV (Thermo Fisher Scientific) plunge freezer with one blotting arm covered by Teflon, excess liquid was removed by blotting the grid between 4-6 s from the back. Vitrification of the sample was achieved by using a mixture of ethane/propane (37%/63%) cooled to liquid nitrogen temperature. Samples were subsequently stored in liquid nitrogen until sample thinning using cryo-focused ion beam milling.

### CryoFIB milling

Cryo-focused ion beam (cryoFIB) milling was performed with the automation methodology previously described (*54*). In brief, vitrified samples were clipped into modified autogrids (Thermo Fisher Scientific) and mounted onto a 40° pre-tilted cryoFIB holder (Leica). The samples were then transferred to an ACE600 (Leica) using a VCT500 (Leica) and sputter coated with a ∼4.5 nm layer of tungsten. Using the VCT500, the holder was subsequently mounted onto a cryo-stage (Leica) inside of a Crossbeam 550 FIB-SEM (Carl Zeiss Microscopy). To prepare the sample for the automated protocol, grids were coated with a layer of organoplatinum. Using the SEM (3 kV, 56 pI) milling targets were identified on the grids and then prepared for the automated milling procedure. During the automation, 12-20 µm lamella were prepared using rough milling (700 pA, 300 pA and 100 pA) and polishing currents (50 pA). We aimed for a final lamella thickness of ∼200-250 nm. To save the milled sample for TEM imaging, grids were unloaded from the Crossbeam 550 using the VCT500 and stored in LN_2_.

### CryoET data collection

Using cryoET, tomographic data was collected of the cryoFIB-milled archaeal lawns. To record the data, we used a Titan Krios equipped with a Gatan K3 direct electron detector (Gatan) and GIF BioQuantum (Gatan, 20 eV slit). To automate tilt series collection, we used SerialEM to collect micrographs within a tilt range of 50° to −70° with 3° dose-symmetric increments. The total dose per tilt series was ∼120-140 e^−^/Å^2^ with ∼10 frames per tilt. The defocus used was ∼8 µm with a pixel size of 3.4 Å.

### Tomogram reconstruction and subtomogram averaging

Collected tilt-series were drift corrected using alignframes and reconstructed with the IMOD package (*55*). All shown tomograms are 4x binned and deconvolution filtered with tom_deconv (*56*). To perform subtomogram averaging, dynamo was used for particle extraction, alignment and averaging. To average the baseplate of the extended *Hb* CIS, an initial set of 68 particles were picked from cryoFIB-milled *Hb* cells. Particles were extracted (88 x 88 x 88-pixel box size) from 4 x 4-binned tomograms (13.6 Å pixel size) and aligned in 6 iterations. To remove poorly aligned particles, the particle dataset was cleaned based on cross-correlation values and aligned for another 6 rounds. The final average has c6 rotational symmetry and had a total of 57 unique extended CIS baseplates particles. The high-resolution atomic structure of the *Anabaena* PCC 7120 (PDB: 7b5h) extended CIS baseplate was fitted into the *Hb* subtomogram average using ChimeraX (*57*). To generate an average of the extended and contracted sheath, 4,522 extended and 3,668 contracted sheath particles were extracted (40 x 40 x 40-pixel box size) from 4 x 4-binned cryoFIB-milled tomograms (13.6 Å pixel size). Particles were aligned over 7 iterations with a final applied rotational symmetry of c6.

### Toxin-immunity gene identification and bioinformatic analysis

The genes identified in the interspacing region of the *Hb* CIS gene cluster (*Hbor_38720-38800*) were each amplified from *Hb* cell lysate and ligated into the expression and C-terminal 6xHis-tagging plasmid - pWL102et. Each generated plasmid was transformed into *Hv* using the transformation methodology previously described. To test if transformations were successful, colonies were tested for the presence of plasmid and insert using PCR. In addition, Western blotting using anti-His antibodies was used to identify if there was expression of the immunity genes from the plasmid. Unsuccessfully transformed plasmids transformations were repeated three times and also transformed into Top10 *E. coli*. To test if the identified toxins were accompanied by immunity genes, the toxin *Hbor_38740* and *Hbor_38760* were amplified together with their downstream partners *Hbor_38750* and *Hbor_38770*, respectively, using PCR. The amplified fragment was then transformed into *Hv.* PCR and Western blotting were again used to verify that identified colonies retained the plasmid with the insert and that the plasmids were expressing. To identify potential functional domains, both toxins were analyzed using InterPro (*58*).

### Identification of haloarcheal CIS gene clusters and phylogenetic analysis

Representative RefSeq genomes belonging to class *Halobacteria* (halophilic archaea) were downloaded from NCBI (*59*). BLASTP (*60*) was used to search these genomes for gene clusters that potentially encode for CISs, by using Cis2 (*Hbor_38820*) as query. The whole *Hb* gene cluster was then searched against these filtered genomes to identify the boundaries of each of the detected CIS gene clusters. These gene clusters were then annotated and visualized using clinker 0.0.28 (*61*). 16S rDNA sequences and Cis11a protein sequences from each strain were extracted for phylogenetics analysis. Sequence alignments were performed through Clustal Omega 1.2.4 (*62*) and the resulting alignments were subsequently trimmed using TrimAl 1.3 (--automated1) (*63*). FastTree 2.1.11 (*64*) and Interactive Tree of Life (*65*) were used to generate and visualize approximately-maximum-likelihood phylogenetic trees using the trimmed alignments as inputs, respectively.

### Bioinformatic analysis of putative membrane anchor

The gene *Hbor_38620* was run through DeepTMHMM to determine if the protein has transmembrane helices and the overall topology of the protein (*66*). The determined transmembrane helices were then identified in the Alphafold structure and displayed using ChimeraX (*57*, *67*).

### Confocal microscopy of putative membrane protein tagged with GFP

A *Hv* strain expressing the GFP-tagged putative membrane protein (*Hbor_38620*) was grown for 24 h at 45°C. A 20 µl aliquot of this culture was spotted onto an agarose pad (2.3 M NaCl, 88.3 mM MgCl_2_·6H_2_O, 85.3 mM MgSO_4_·7H_2_O, 56.5 mM KCl, 12 mM Tris.HCl pH 7.5, 3 mM CaCl_2_ 1% agarose) and left to dry at room temperature. After 30 min, a cover slip was placed onto the sample and taken to be imaged using the Zeiss LSM900 with Airyscan 2 detector and a x63 / 1.4 NA oil-immersion objective. *z*-stacks of target cells were taken using a confocal imaging track detecting transmitted light and a separate Airyscan track detecting GFP. Collected confocal stacks were deconvolved with the Zeiss LSM Plus processing function and the Airyscan images were processed using Zeiss deconvolution (jDCV, 10 iterations) in ZenBlue (v.3.5). Minimum-intensity projections of the transmitted light channel and extraction of single Airyscan slices were performed in Fiji (*53*).

### Statistical Analysis

Data analysis and statistics mentioned in the text, including standard deviations and 2way ANOVA analysis, were performed in Microsoft Excel and GraphPad Prism.

## Acknowledgments

ScopeM is acknowledged for providing instrument access at ETH. The Functional Genomics Center Zurich is acknowledged for collecting and analyzing mass spectroscopy data and providing technical expertise. We would like to acknowledge Florian Wollweber for assisting with light microscopy imaging. We thank Hanna Wetzel for cloning pSVA5921.

## Funding

Swiss National Science Foundation (310030_212592) (MP)

European Research Council (679209/101000232) (MP)

NOMIS foundation (MP)

Deutsche Forschungsgemeinschaft (403222702-SFB 138) (SS, SVA)

VW Momentum grant (Grant number 94933) (PN)

## Author contributions

TZ conducted all experiments that involved genetics, sheath preps, negative stain electron microscopy, grid preparation for electron microscopy, building and validating atomic models, growth curves, functional assays, cryoET sample preparation/imaging, tomogram reconstruction and subtomogram averaging. JJM and TZ performed the bioinformatic analysis. AA, PN and SS assisted with establishing the time lapse live cell imaging of halophiles. JX processed the single particle dataset. SVA provided supervision to PN and SS and provided support in archaeal genetics and physiology. TZ and MP wrote the manuscript with input from all coauthors.

## Data and material availability

Representative tomograms and subtomogram averages will be submitted to EMDB (accession #XXX). CryoEM structures will be submitted to PDB (accession #XXX).

## Supplementary Figures

**Figure S1:**
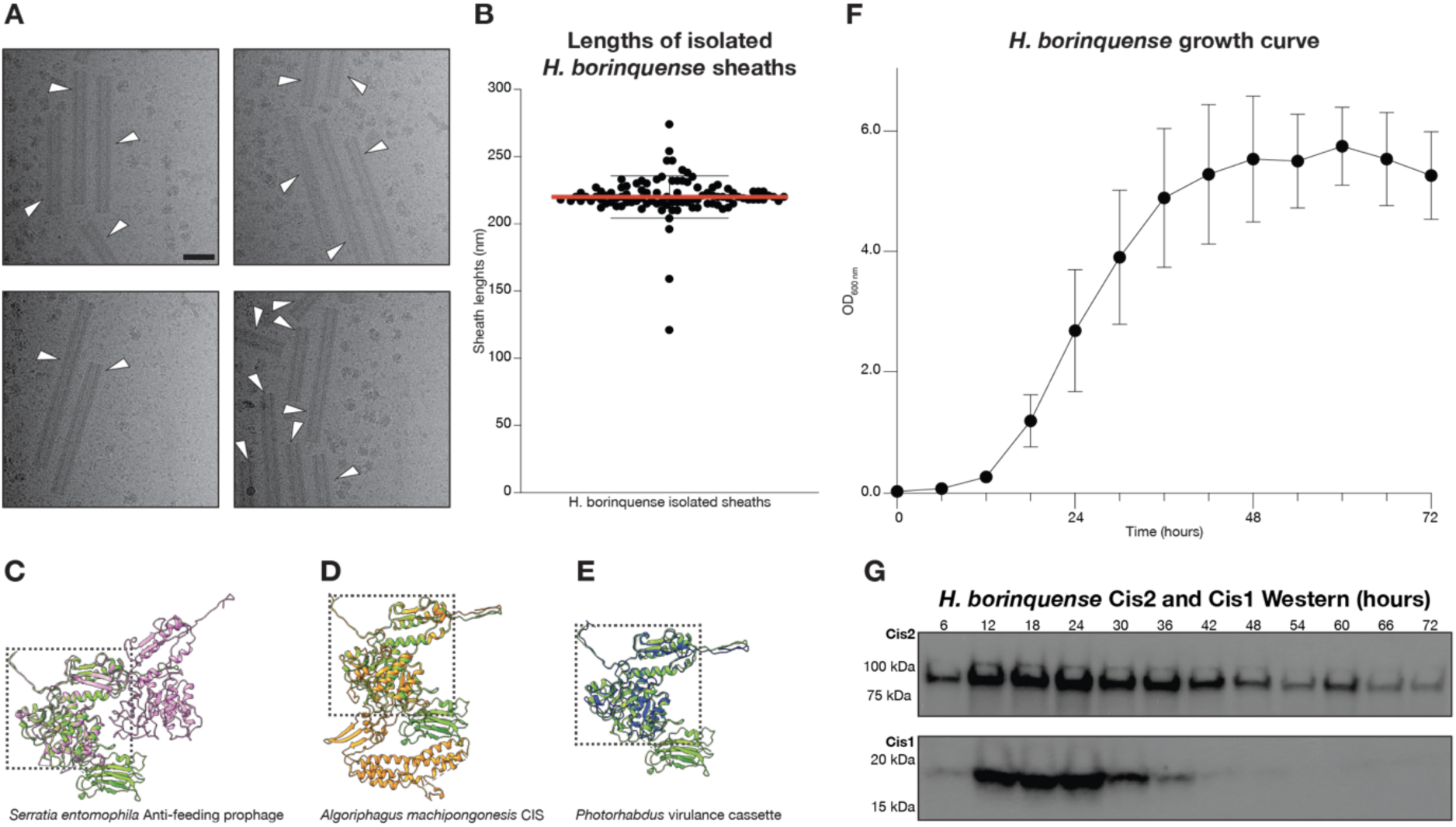
Sheath length distribution, structure comparison and Western blot analysis. **(A)** Four example EM micrographs of sheath purifications, revealing length variations of contracted *H. borinquense* sheaths (arrowheads). Scale bar: 50 nm. **(B)** Chart showing the length distribution of 100 sheath from 89 micrographs. A mean length of 220 nm ±16 nm SD was measured with a maximum length of 274 nm and a minimum length of 121 nm. **(C-E)** Superposition and alignment of the contracted sheath monomer from *H. borinquense* onto that of *Serratia entomophila* in pink (PDB: 6RC8, (*21*)), *Algoriphagus machipongonesis* in orange (PDB: 7AEK, (*24*)) and the *Photorhabdus* virulence cassette in blue (PDB: 6J0C, (*22*)). The structures show high structural similarities in domains 1 and 2 (outlined by the black box) with large variations in domain 3. **(F)** Growth curve of a *H. borinquense* culture initiated at OD 0.01 with a late exponential phase pre-culture. *H. borinquense* was grown in 25 ml YPC at 45°C and 200 rpm. Points were measured every 6 h over the course of 72 h. **(G)** Western blot of *H. borinquense* samples at 6 h intervals. Using antibodies, against the Cis2 (sheath, ∼63.3 kDa) and Cis1 (inner tube, ∼16.4 kDa), the expression levels of these two proteins was analyzed over 72 h. Samples loaded on the gel were normalized to be at OD 1.0. Expression of sheath remained similar during exponential phase, however, Cis1 expression dropped at mid-exponential phase around 24 h.

**Figure S2:**
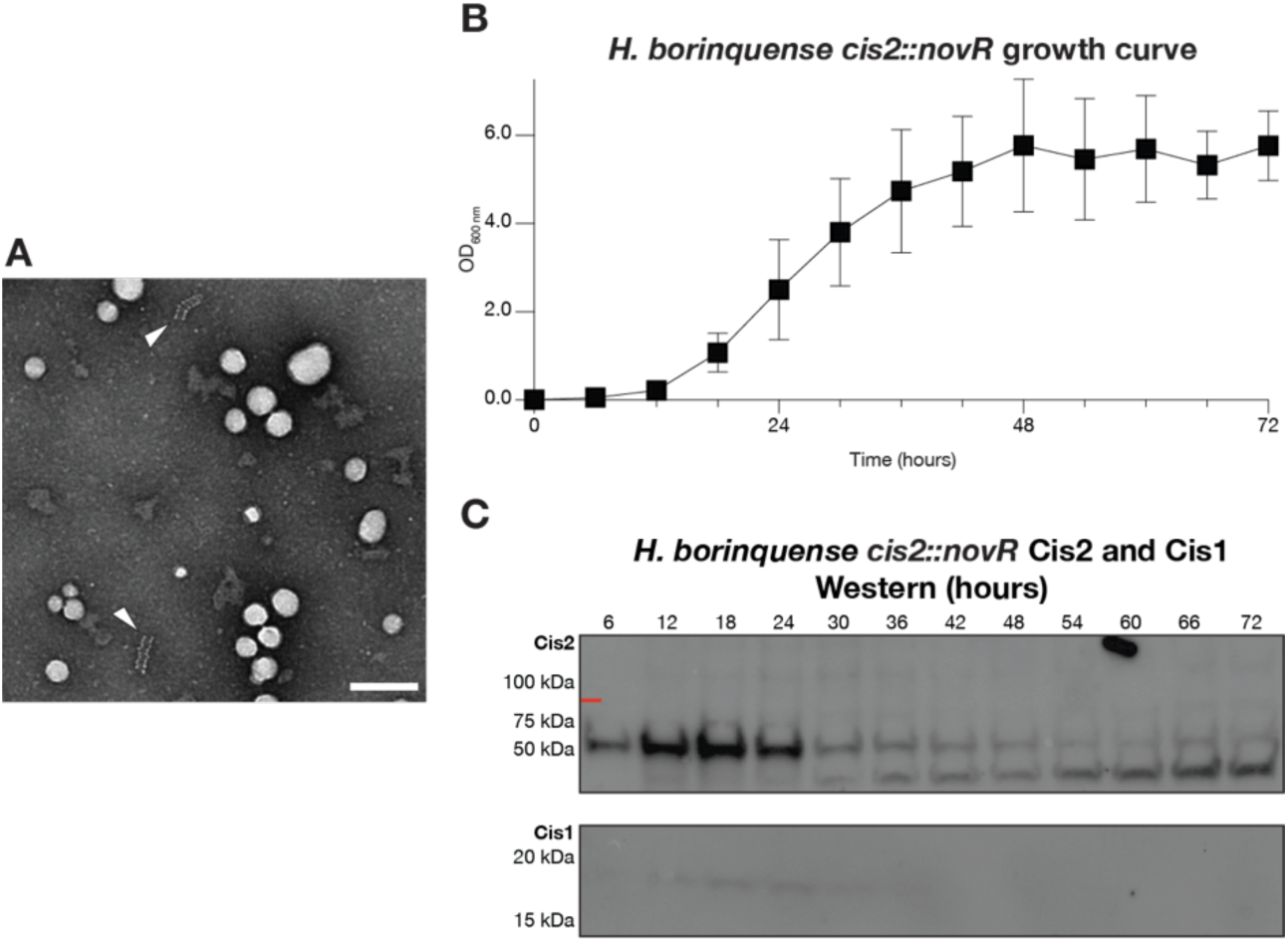
*H. borinquense* c*is2*::*novR* mutant analysis. **(A)** Representative negative stain EM micrograph of the sheath isolates from the c*is2*::*novR* mutant strain generated by homologous recombination. Few potential sheath-like structures were observable. Identified structures are shorter and look irregular (white arrow). Scale bar: 200 nm. **(B)** Growth curve of a *H. borinquense* c*is2*::*novR* culture initiated at OD 0.01 with a late exponential phase pre-culture. The *H. borinquense* sheath mutant was grown in 25 ml YPC at 45°C and 200 rpm. Points were measured every 6 h over the course of 72 h. **(C)** Western blot of *H. borinquense* c*is2*::*novR* growth curve collected at 6 h intervals. Using antibodies against Cis2 (sheath, ∼63.3 kDa) and Cis1 (inner tube, ∼16.4 kDa), the expression levels of these two proteins was analyzed over 72 h. Samples loaded on the gel were normalized to be at OD 1.0. Bands for the Cis2 correspond to a much shorter protein than that of the full-length Cis2 (red line) and Cis1 expression was barely visible in blots.

**Figure S3:**
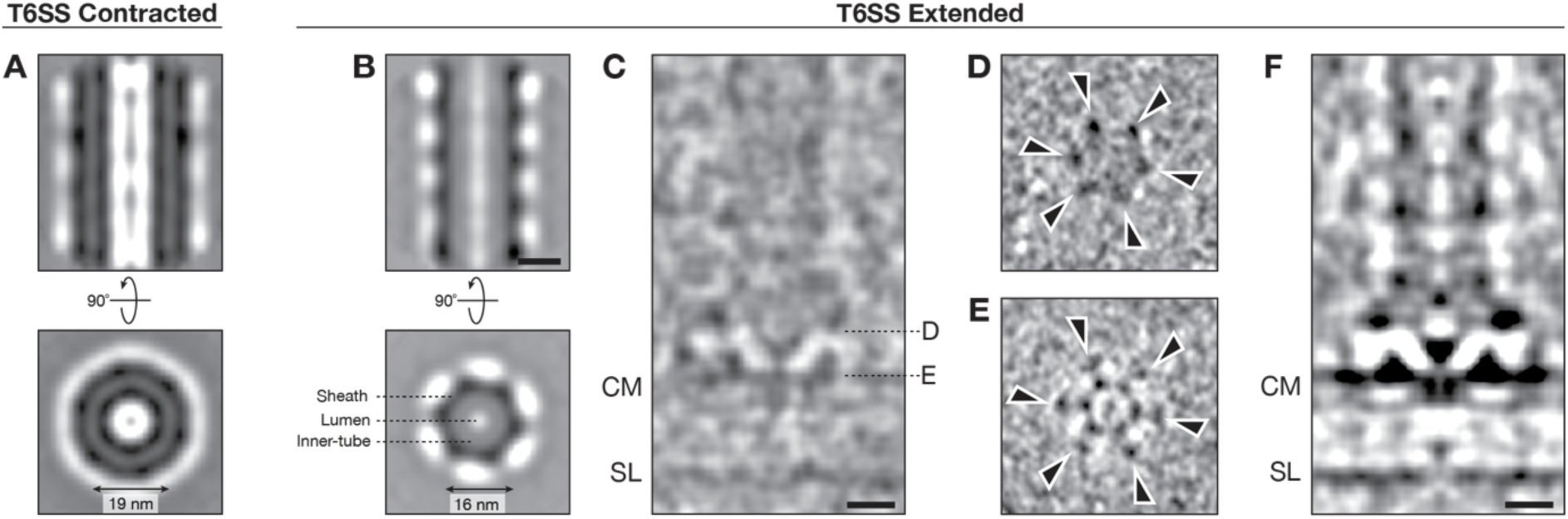
The T6SS is composed of sheath-tube, baseplate and membrane-anchor modules that possess C6 symmetry. **(A)** Contracted T6SS sheath subtomogram average. Shown are 1.36 nm thick longitudinal and perpendicular slices through the subtomogram average. In the contracted state, the sheath has a diameter of around 19 nm with no clear density in the center. **(B)** Subtomogram average of the T6SS sheath in the extended state. Shown are 1.36 nm thick longitudinal and perpendicular slices through the subtomogram average. In the extended state, the sheath-tube module has a diameter of around 16 nm. A clear density for the sheath is observable with a more diffuse density for the inner tube. In the subtomogram average the lumen of the inner tube appears to be empty. **(C-E)** Subtomogram average of the *H. borinquense* T6SS in the extended state without any applied symmetry. The average consists of 57 particles extracted from *H. borinquense* tomograms. Shown are 1.36 nm thick longitudinal and perpendicular slices. In panel **C** all three modules (sheath-tube, baseplate and membrane anchor) of the *H. borinquense* T6SS are visible. Panel **D** is a slice through the baseplate and shows six densities at the edge of the baseplate (arrowheads). Panel **E** is a slice just above the cytoplasmic membrane and shows that there are 13 membrane attachment points. Six contacts are made by both the inner and outer ring (‘feet-like’ structures highlighted using arrowheads), and a single attachment at the center of structure (‘cage’). Scale bar: 10 nm. **(F)** Average in panel **A** with an applied C6 rotational symmetry. Scale bar: 10 nm.

**Figure S4:**
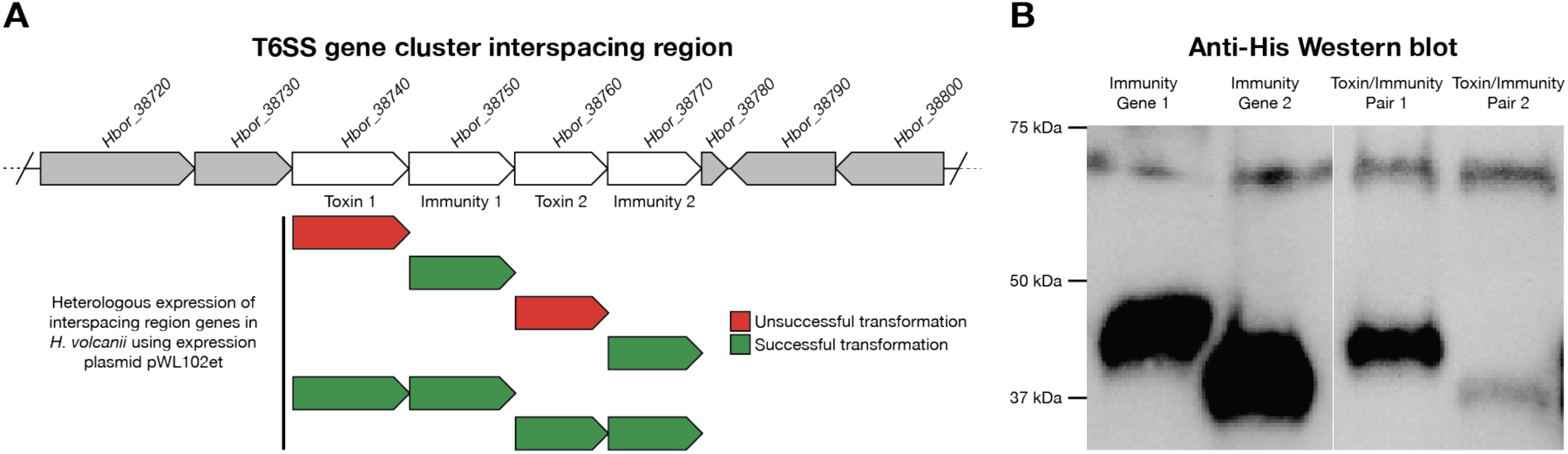
The interspacing region of the T6SS gene cluster contains two putative toxin-immunity pairs. **(A)** Schematic of the interspacing region of the *H. borinquense* T6SS gene cluster. Depicted is transformation success of genes or gene pairs in the interspacing region when transformed into *H. volcanii* using the pWL102et expression plasmid. **(B)** Anti-His Western blotting of *H. volcanii* transformed with plasmids shown in **A**. The expression of the immunity genes (immunity 1 = 38.9 kDa, immunity 2 = 30.5 kDa) was detectable in both individual expression of the gene and during expression in tandem with the toxin pair.

**Figure S5:**
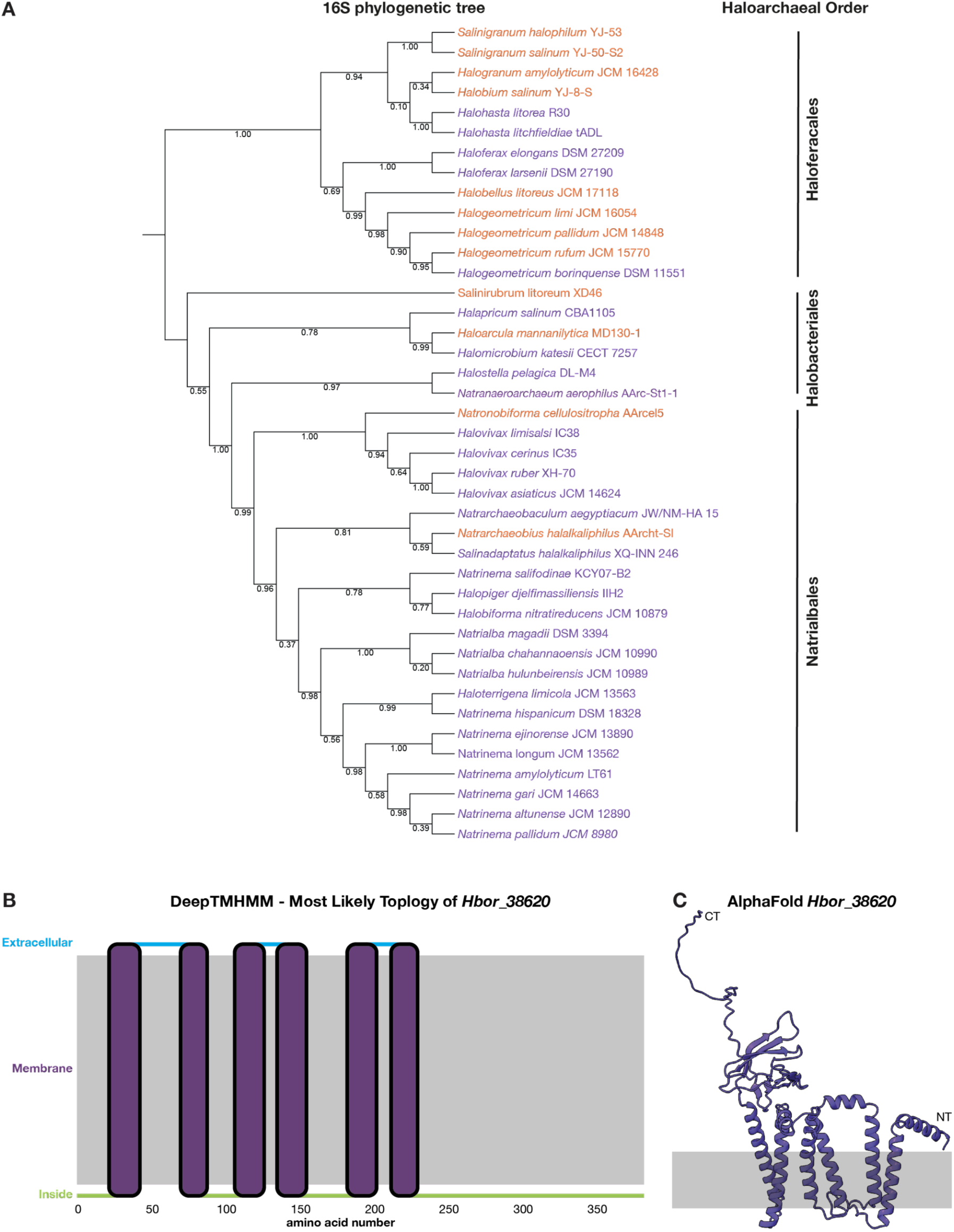
Bioinformatic analysis of haloarchaea CIS gene clusters reveals conserved putative membrane anchor. **(A)** 16S rDNA phylogenetic tree of haloarchaeal strains which have a CIS gene cluster. Representatives of three out of four different haloarchaeal orders can be identified. Color coding of the species matches Figure 4A. The mismatch between 16S rDNA and Cis11a phylogenies indicates horizontal gene transfer of the CIS gene clusters. **(B)** Results of the DeepTMHMM transmembrane helices prediction tool indicates that *Hbor_38620* has six transmembrane helices. It also predicts that the putative membrane protein has both the C-and N-terminus localized in the cytoplasm of the cell. **(C)** AlphaFold-predicted structure of *Hbor_38620* embedded within the cytoplasmic membrane. The putative membrane protein is predicted to have six transmembrane helices with a highly unstructured C-terminus. We speculate that this C-terminus may facilitate T6SS membrane anchoring by interacting with T6SS baseplate components.

**Figure S6:**
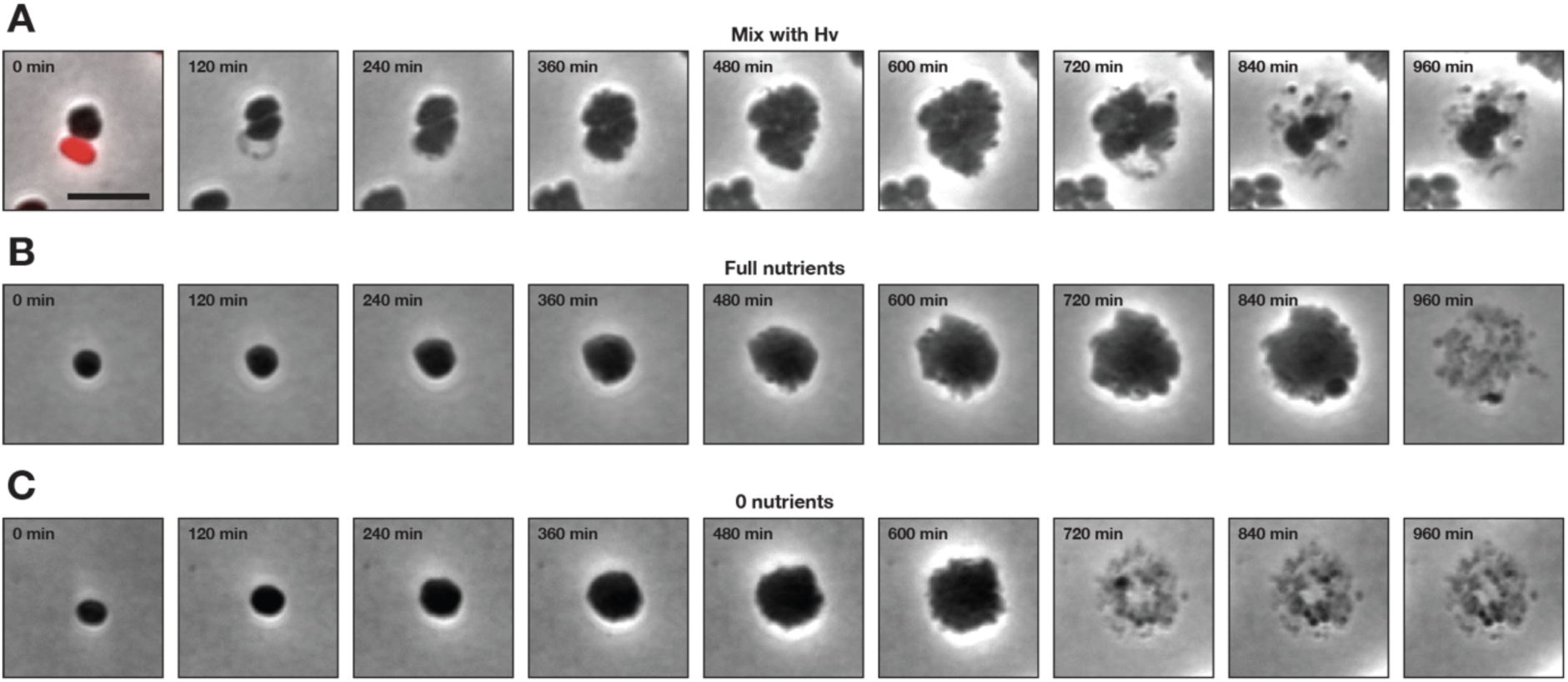
*H. elongans* occasionally undergoes cell lysis during time lapse imaging. **(A-C)** Time lapse imaging (15 min frame rate) of *H. elongans* cells that undergo cell lysis while imaged over 960 min at 45°C. In panel **A**, a selection of frames are shown as overlays of phase contrast images with red fluorescence (respective time points are indicated). Red fluorescence originated from mScarlet expressed in *H. volcanii*. During imaging, *H. volcanii* is observed to undergo a “cell lysis” event at a time point between 0 and 120 min. *H. elongans* is also observed to begin lysing between time point 600 and 960 min. In panel **B** and **C**, a selection of frames are shown in which *H. elongans* grows in the absence of *H. volcanii*. Both under full and 0 nutrient conditions *H. elongans* undergoes “cell lysis” before the end of the time lapse imaging. Scale bar: 5 µm.

**Table S1:**
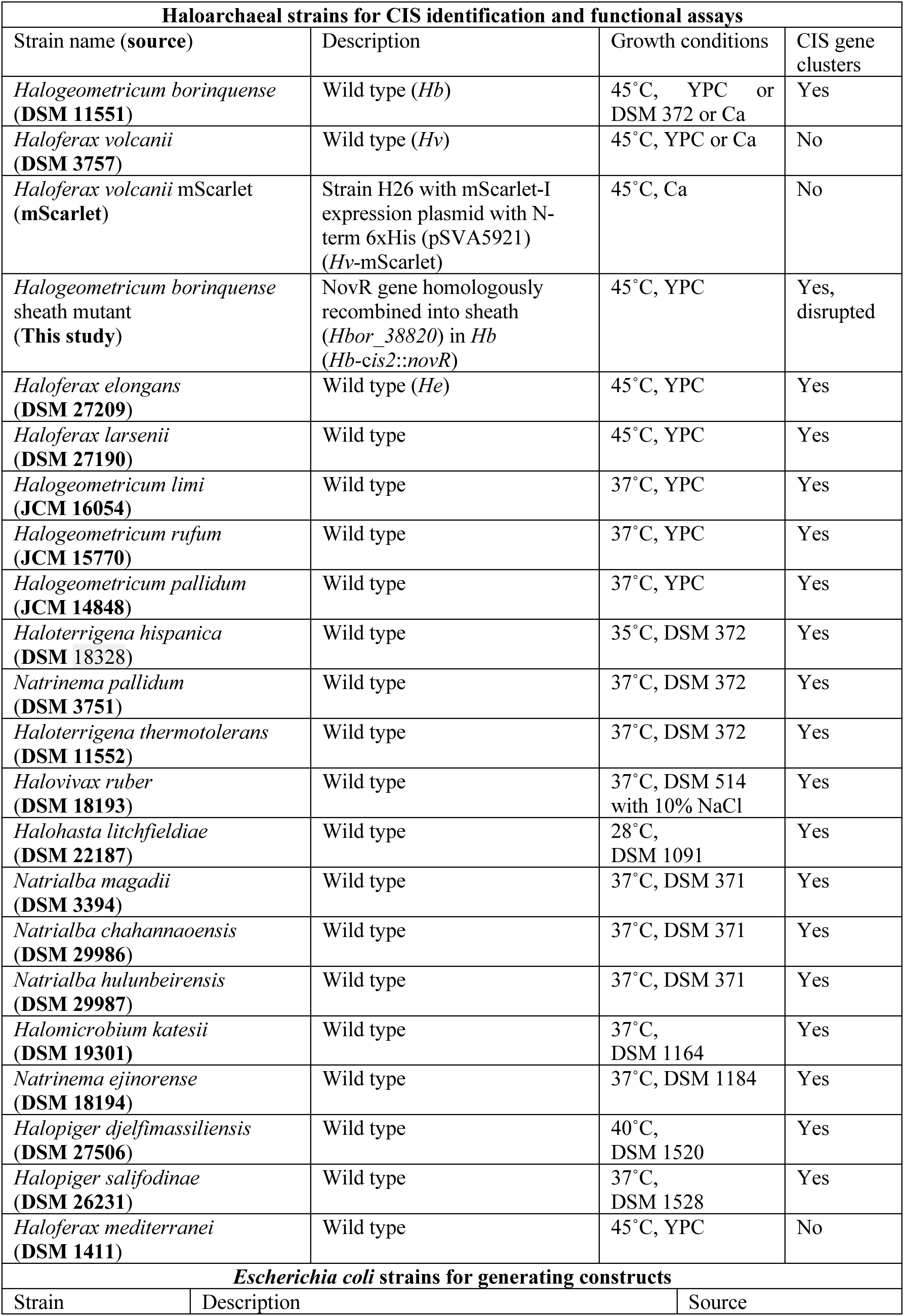
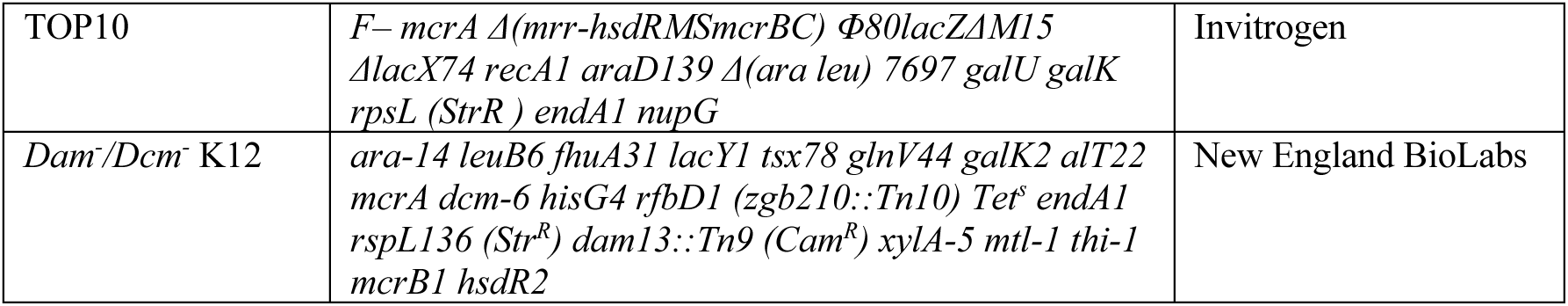
Strains and culturing conditions.

| **Haloarchaeal strains for CIS identification and functional assays** |  |  |  |
| --- | --- | --- | --- |
| Strain name (**source**) | Description | Growth conditions | CIS gene clusters |
| *Halogeometricum borinquense* (**DSM 11551**) | Wild type (*Hb*) | 45°C, YPC or DSM 372 or Ca | Yes |
| *Haloferax volcanii* (**DSM 3757**) | Wild type (*Hv*) | 45°C, YPC or Ca | No |
| *Haloferax volcanii* mScarlet (**mScarlet**) | Strain H26 with mScarlet-I expression plasmid with N-term 6xHis (pSVA5921) (*Hv*-mScarlet) | 45°C, Ca | No |
| *Halogeometricum borinquense* sheath mutant (**This study**) | NovR gene homologously recombined into sheath (*Hbor_38820*) in *Hb* (*Hb-cis2::novR*) | 45°C, YPC | Yes, disrupted |
| *Haloferax elongans* (**DSM 27209**) | Wild type (*He*) | 45°C, YPC | Yes |
| *Haloferax larsenii* (**DSM 27190**) | Wild type | 45°C, YPC | Yes |
| *Halogeometricum limi* (**JCM 16054**) | Wild type | 37°C, YPC | Yes |
| *Halogeometricum rufum* (**JCM 15770**) | Wild type | 37°C, YPC | Yes |
| *Halogeometricum pallidum* (**JCM 14848**) | Wild type | 37°C, YPC | Yes |
| *Haloterrigena hispanica* (**DSM 18328**) | Wild type | 35°C, DSM 372 | Yes |
| *Natrinema pallidum* (**DSM 3751**) | Wild type | 37°C, DSM 372 | Yes |
| *Haloterrigena thermotolerans* (**DSM 11552**) | Wild type | 37°C, DSM 372 | Yes |
| *Halovivax ruber* (**DSM 18193**) | Wild type | 37°C, DSM 514 with 10% NaCl | Yes |
| *Halohasta litchfieldiae* (**DSM 22187**) | Wild type | 28°C, DSM 1091 | Yes |
| *Natrialba magadii* (**DSM 3394**) | Wild type | 37°C, DSM 371 | Yes |
| *Natrialba chahannaoensis* (**DSM 29986**) | Wild type | 37°C, DSM 371 | Yes |
| *Natrialba hulunbeirensis* (**DSM 29987**) | Wild type | 37°C, DSM 371 | Yes |
| *Halomicrobium katesii* (**DSM 19301**) | Wild type | 37°C, DSM 1164 | Yes |
| *Natrinema ejinorensis* (**DSM 18194**) | Wild type | 37°C, DSM 1184 | Yes |
| *Halopiger djelfmassiliensis* (**DSM 27506**) | Wild type | 40°C, DSM 1520 | Yes |
| *Halopiger salifodinae* (**DSM 26231**) | Wild type | 37°C, DSM 1528 | Yes |
| *Haloferax mediterranei* (**DSM 1411**) | Wild type | 45°C, YPC | No |
| **Escherichia coli strains for generating constructs** |  |  |  |
| Strain | Description | Source |  |

|  |  |  |
| --- | --- | --- |
| TOP10 | *F– mcrA Δ(mrr-hsdRMSmcrBC) Φ80lacZΔM15 ΔlacX74 recA1 araD139 Δ(ara leu) 7697 galU galK rpsL (Str<sup>R</sup>) endA1 nupG* | Invitrogen |
| *Dam<sup>-</sup>/Dcm<sup>-</sup>* K12 | *ara-14 leuB6 fhuA31 lacY1 tsx78 glnV44 galK2 alt22 mcrA dcm-6 hisG4 rfbD1 (zgb210::Tn10) Tet<sup>s</sup> endA1 rspL136 (Str<sup>R</sup>) dam13::Tn9 (Cam<sup>R</sup>) xylA-5 mtl-1 thi-1 mcrB1 hsdR2* | New England BioLabs |

**Table S2:** Plasmids used in this study.

| Plasmid (source) | Description | Application |
| --- | --- | --- |
| pWL102 (DSM 5717) | *E. coli* - *Hv* shuttle vector for selection and maintenance in both bacteria and archaea; ampicillin and lovastatin resistance | Vector used to generate expression plasmids and in functional assays |
| pWL102et (This study) | Plasmid with pWL102 backbone, Pr R promoter (68) and 6xHis-tag | For expressing and tagging proteins in *Hv* |
| pGB68 (69) | *E. coli* - *Hv* shuttle vector for selection and maintenance in both bacteria and archaea; ampicillin and novobiocin resistance | Used to amplify NovR gene |
| pWL102et NovR (This study) | Plasmid with pWL102et backbone and NovR from pGB68 | Expression of the antibiotic novobiocin resistance gene |
| pWL102et int1 (This study) | Plasmid with pWL102et backbone and *Hbor_38720* from the *Hb* genome | Expression of interspacing gene 1 |
| pWL102et int2 (This study) | Plasmid with pWL102et backbone and *Hbor_38730* from the *Hb* genome | Expression of interspacing gene 2 |
| pWL102et int3 (This study) | Plasmid with pWL102et backbone and *Hbor_38740* from the *Hb* genome | Expression of interspacing gene 3 |
| pWL102et int4 (This study) | Plasmid with pWL102et backbone and *Hbor_38750* from the *Hb* genome | Expression of interspacing gene 4 |
| pWL102et int5 (This study) | Plasmid with pWL102et backbone and *Hbor_38760* from the *Hb* genome | Expression of interspacing gene 5 |
| pWL102et int6 (This study) | Plasmid with pWL102et backbone and *Hbor_38770* from the *Hb* genome | Expression of interspacing gene 6 |
| pWL102et int7 (This study) | Plasmid with pWL102et backbone and *Hbor_38780* from the *Hb* genome | Expression of interspacing gene 7 |
| pWL102et int8 (This study) | Plasmid with pWL102et backbone and *Hbor_38790* from the *Hb* genome | Expression of interspacing gene 8 |
| pWL102et int9 (This study) | Plasmid with pWL102et backbone and *Hbor_38800* from the *Hb* genome | Expression of interspacing gene 9 |
| pWL102et int3/4 (This study) | Plasmid with pWL102et backbone and *Hbor_38740/38750* from the *Hb* genome | Expression of the immunity/toxin pair 1 |
| pWL102et int5/6 (This study) | Plasmid with pWL102et backbone and *Hbor_38760/38770* from the *Hb* genome | Expression of the immunity/toxin pair 2 |
| pET15b (Novagen) | Cloning/expresson plasmid with ampicillin resistance | Vector used to make sheath insertion plasmid |
| pET15b NovR sheath ins (This study) | Plasmid with pET15b backbone and sheath insertion fragment | Plasmid for generating *Hb* sheath insertion mutant through homologous recombination |
| pIDJL40 (70) | Plasmid which provided halotolerant GFP | Used to amplify GFP sequence |
| pWL102et *Hbor_38620*-GFP (This study) | Plasmid with pWL102et backbone and *Hbor_38620* from the *Hb* genome with a GFP tag | Expression of putative membrane protein with C-terminal GFP tag |

**Table S3:**
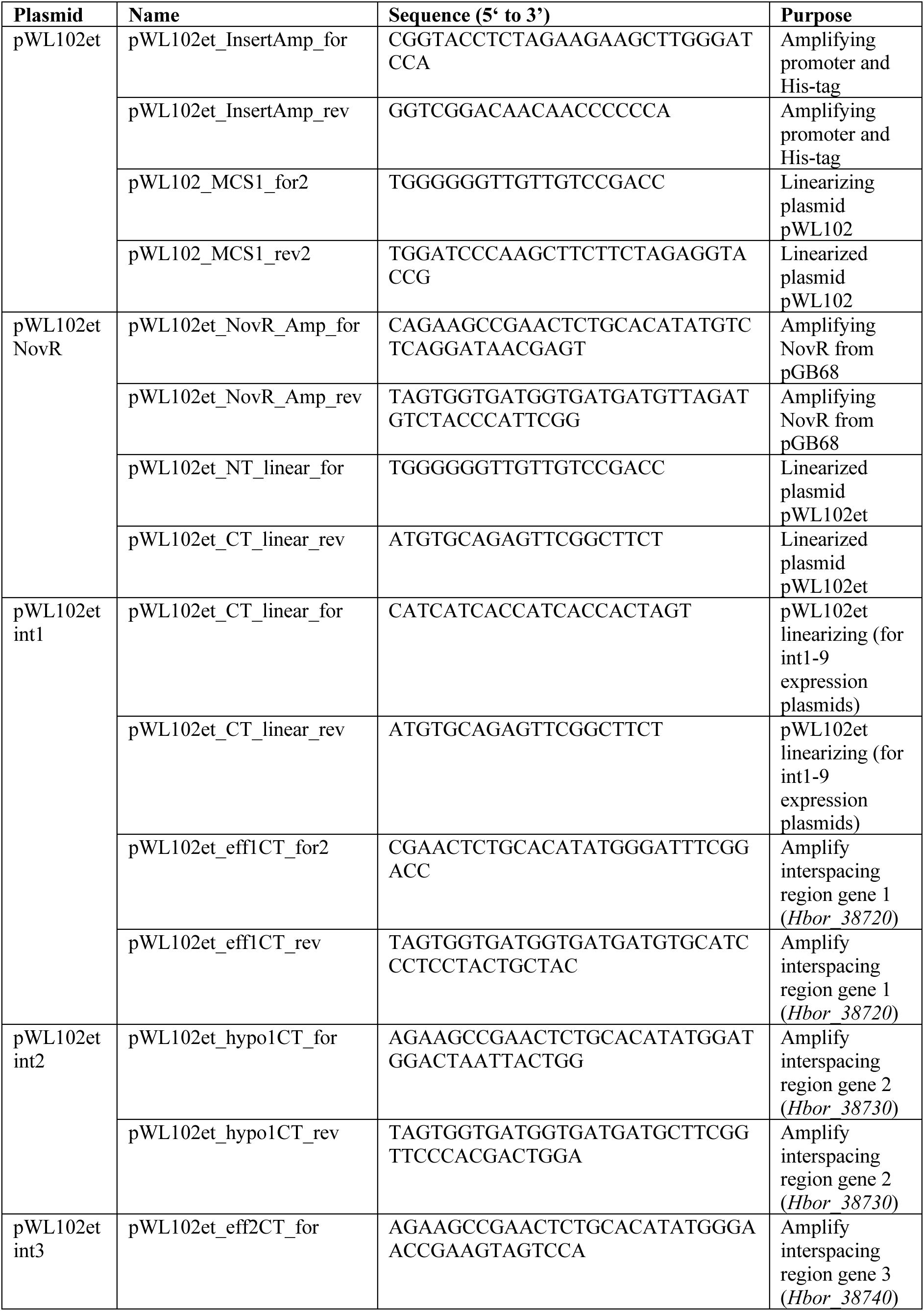
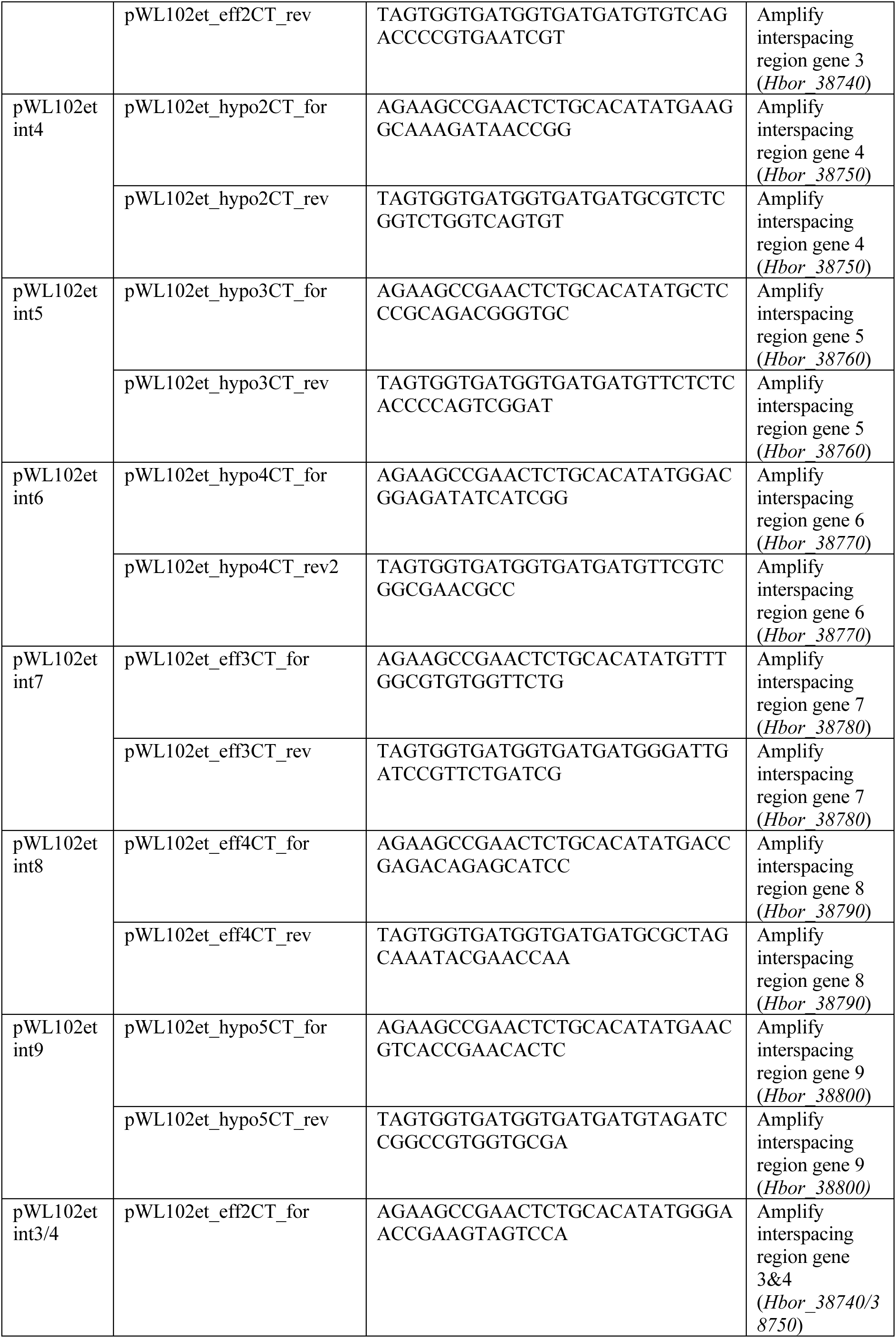
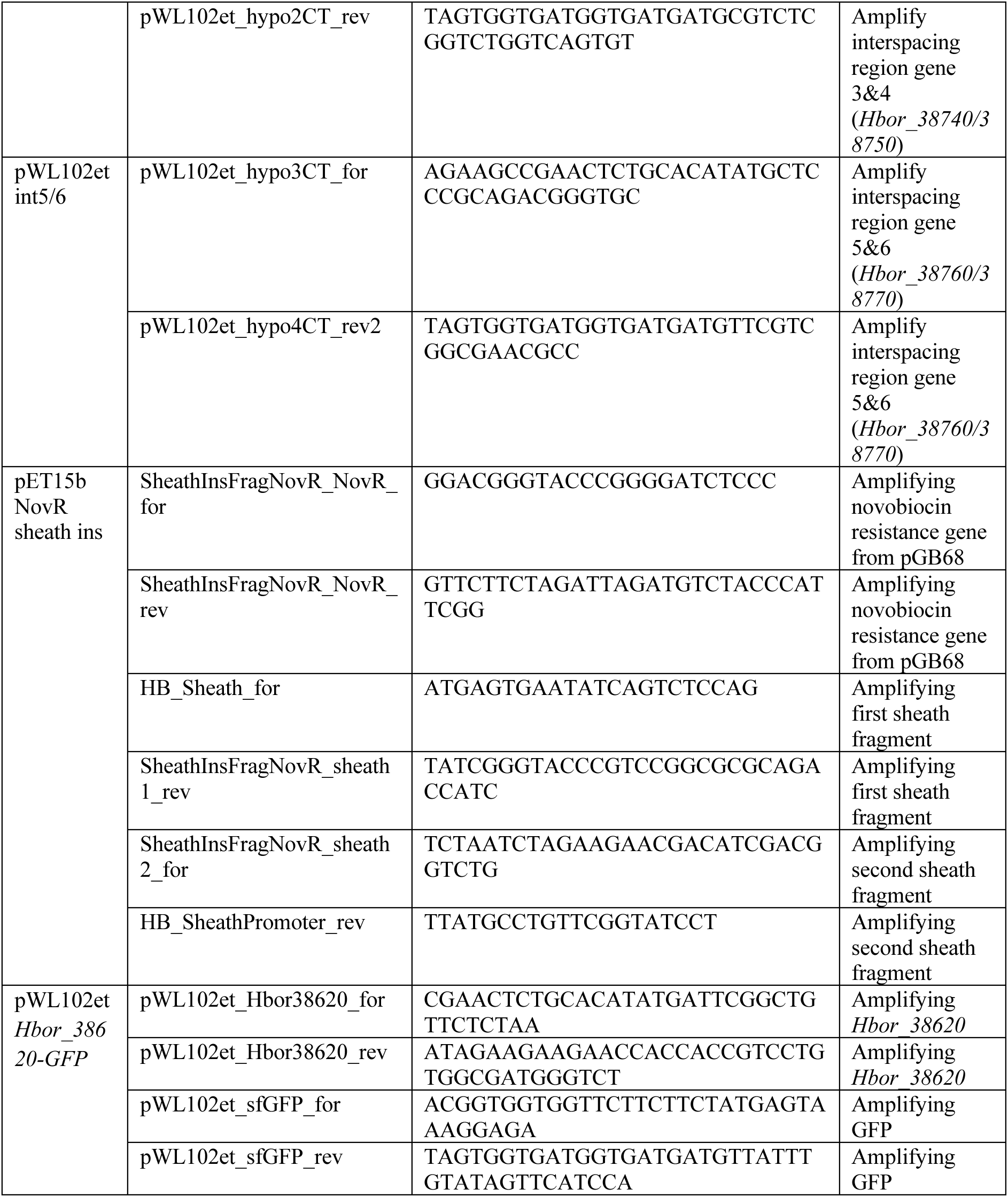
Oligonucleotides used in this study.

**Table S4:** Structure validation of contractile sheath.

| Data collection and processing |  |
| --- | --- |
| Nominal magnification | 81,000 |
| Voltage (kV) | 300 |
| Electron exposure (e<sup>-</sup>/Å) | ~65 |
| Defocus range (μm) | -1 to -3.3 |
| Pixel size (Å/pixel) | 1.10 |
| Symmetry imposed | C6 + helical<br>(twist = 33.24°, rise = 17.09 Å) |
| Final particles (No.) | 127,891 |
| Map resolution | 2.9 |
| FSC threshold | 0.143 |
| Refinement |  |
| Map sharpening B factor (Å<sup>2</sup>) | -86.8 |
| Model composition |  |
| Non-hydrogen atoms | 68,022 |
| Protein residues | 8,766 |
| Chains | 18 |
| R.M.S deviations |  |
| Bond length (Å) | 0.004 |
| Bond angles (°) | 0.708 |
| Validation |  |
| MolProbity score | 1.09 |
| Clashscore | 2.95 |
| Rotamer outlier (%) | 0.00 |
| Ramachandran plot |  |
| Favored (%) | 98.34 |
| Allowed (%) | 1.66 |
| Outlier (%) | 0.00 |
| Masked CC | 0.87 |

**Movie S1: *H. volcanii* “cell lysis” event upon *H. borinquense* contact.**

Six time lapse imaging examples of *H. volcanii* “lysis” events when in direct contact with *H. borinquense*. Red fluorescence signal is used identify mScarlet expressing *H. volcanii* from *H. borinquense*. Scale bar: 5 µm.

**Movie S2: *H. volcanii* “no proliferation” event upon *H. borinquense* contact.**

Six time lapse imaging examples of *H. volcanii* “no proliferation” events when in direct contact with *H. borinquense*. Red fluorescence signal is used identify mScarlet expressing *H. volcanii* from *H. borinquense*. Scale bar: 5 µm.

